# Young Asian elephant calves show differentiated social relationships with conspecifics

**DOI:** 10.1101/2025.04.27.650831

**Authors:** T. Revathe, T.N.C. Vidya

## Abstract

Social relationships may increase longevity and reproductive success of individuals, thereby providing crucial benefits in social species. Adults of many social species have differentiated social relationships; however, when and how these relationships form and are maintained are often not understood. We studied spatial and behavioural interactions between young calves (≤6 months old) and conspecific females (≥5 years old) to understand social ontogeny in a wild Asian elephant (*Elephas maximus*) population in Nagarahole and Bandipur National Parks, southern India. We found that elephant groups with calves comprised females – mother and ‘escort’, who provided active care, and females who did not provide overt, active care – ‘other females’. Escorts, apart from coordinating their movement with the calf, provided allomaternal care by initiating positive and helpful interactions with the calf. Other females rarely initiated interactions, which were usually negative, towards calves. Primarily calves, rather than conspecific females, initiated and terminated proximity contacts and behavioural interactions. Calves were near and interacted similarly with their mothers and escorts, under feeding, resting, and social contexts, and almost never interacted with other females. Similarly, we found remarkable similarities between mothers and escorts in their behaviours towards calves, with the primary difference being that escorts did not provide milk. Our results suggest that calves begin to form differentiated social relationships from a young age. The presence of calves and their interactions with escorts might result from existing close social relationship between mothers and escorts or may establish new relationships. Allomaternal care may also reinforce relationship differentiation in such social species. As calf-escort interactions involved behaviours that may potentially be helpful in protection and feeding skill acquisition, the development and survival of calves may be affected by the presence of, and interactions, with allomothers.

**Graphical Abstract:** 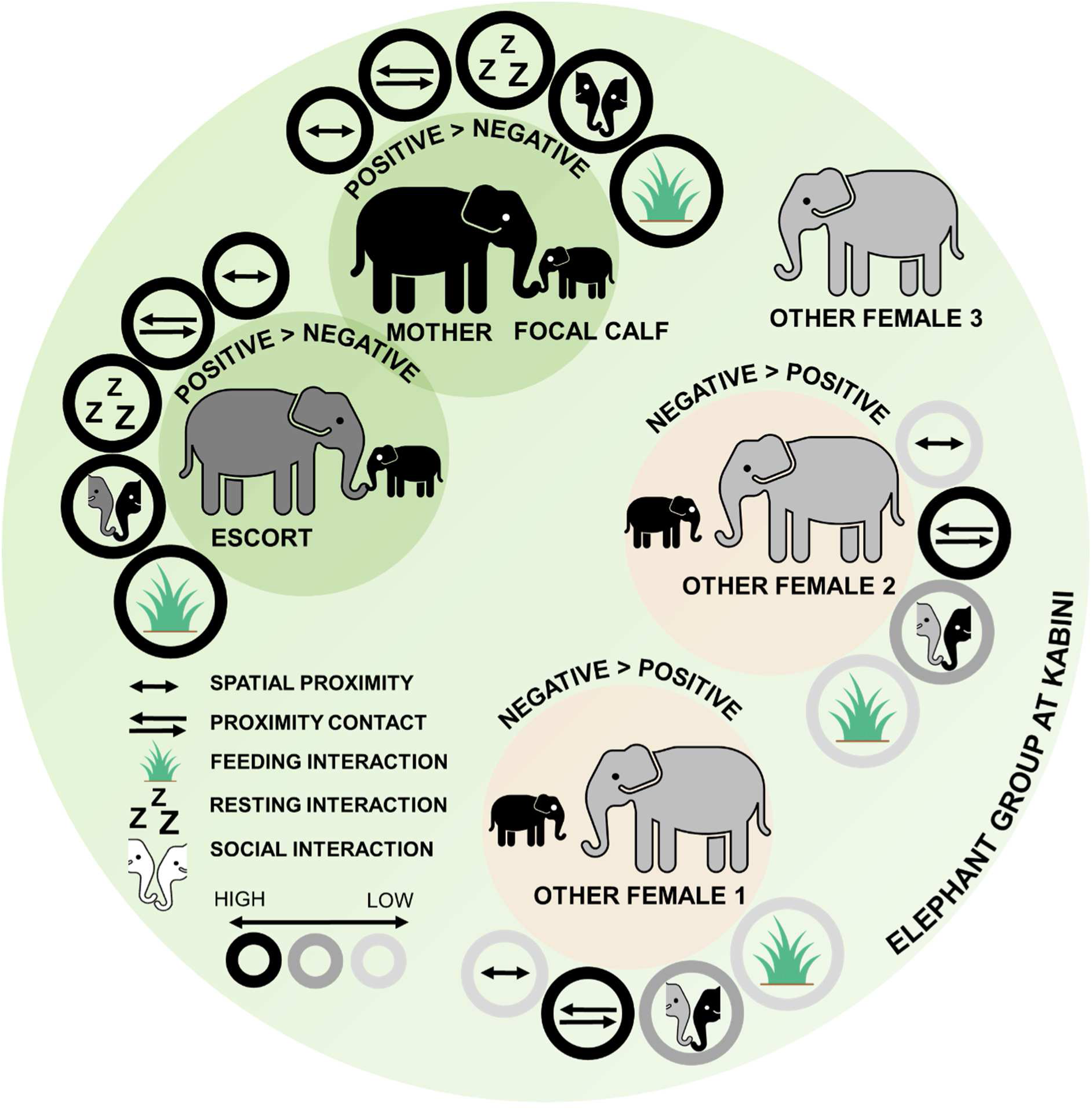

We studied spatial and behavioural interactions between young calves (≤6 months old) and conspecific females to understand social ontogeny in a wild Asian elephant population. We found differentiated relationships between calves and conspecifics even at a young age. Escorts, who showed coordinated movement with calves, engaged in other positive interactions with calves, with remarkable similarity to the mothers, whereas other females rarely initiated such interactions. Escorts are thus allomothers in the Kabini elephant population. Calves, rather than conspecifics, initiated and terminated most of the proximity and behavioural interactions and were thus precocial in forming new relationships.

## Introduction

The development of social relationships is a central component of the ontogeny of social mammals (Bekoff, 1972). Differentiated social relationships may be beneficial (Connor et al., 2001; Langergraber et al., 2013), and such relationships may exist from a young age itself or form gradually (Berman, 1982a, 1982b; Berman et al., 1997; de Waal, 1990; Roatti et al., 2023). Social relationships involving care are of particular importance to developing individuals. In mammals, young individuals form their closest relationship with their mother, who provides a majority of the care (van Noordwijk, 2012). In some species, parental care may be supplemented by alloparental care or allocare, which refers to care contributed by conspecifics apart from the biological parent(s) in raising offspring that are not one’s own.

Alloparental care is documented in over 100 social mammals and include behaviours such as babysitting or accompanying the young one, carrying, active food provisioning, allosuckling, and defending against conspecifics or predators that result in direct interactions between allocarers and the young one (Brotherton et al., 2001; Clutton-Brock et al., 2000; Erb & Porter, 2020; Estes & Goddard, 1967; Gadgil & Nair, 1984; Gero et al., 2013; Hill & Campbell, 2014; Hodge, 2005; Hrdy, 1976; Lee, 1987; Pal et al., 2021; Pusey & Packer, 1994; Rood, 1978; Rowell et al., 1964; Whitehead, 1996; Xiang et al., 2019). Allocare may be provided by one or more or all the members of a social group, and allocarers may be generalised or may specialise in caretaking tasks (Erb & Porter, 2020; Garber, 1997; Gero et al., 2009). Young ones may solicit help from specific allocarers and/or allocarers may approach specific young ones to initiate affiliative interactions (Brotherton et al., 2001; Gilchrist, 2004). Ultimately, allocarers can interact differently than other conspecifics with the young ones and *vice versa*, resulting in the development of differentiated social relationships at a young age.

Among social mammals, primates are known for their highly differentiated social relationships (Berman, 1982a, 1982b; Roatti et al., 2023; Silk et al., 2009, 2010). Like many primates, adult female elephants show complex and differentiated social structure (Archie et al., 2011; de Silva et al. 2011; Nandini et al., 2017, 2018; Wittemyer et al., 2005), but when and how relationships form and become differentiated are not well studied. Elephants live in matrilineal societies with female philopatry and male dispersal (Archie et al., 2005; Vidya & Sukumar, 2005). Males associate only temporarily with female groups after dispersal (Keerthipriya et al., 2021) and do not provide parental care (McKay, 1973). Female elephants give birth to a single calf at a time; twins are very rare. Calves are precocial at birth but are nutritionally and socially dependent on their mothers (Gadgil & Nair, 1984; Lee & Moss, 2011; Nair, 1989; Revathe et al., 2020). Young individuals (≤2.5 years of age) face high predation risk from lions in Africa (Loveridge et al., 2006) and tigers in India (Williams, 1950). Young ones are weaned at only 4-5 years of age or after the birth of a sibling (Mar et al., 2012) and constantly associate with the mother until then. Thus, female elephants invest heavily in their young for an extended duration, matched or exceeded only by certain great apes and cetaceans (Mann, 2019; van Noordwijk et al., 2018).

Allomaternal care has been conjectured to be important in the evolution and cohesiveness of female elephant societies (Dublin, 1983; Gadgil & Nair, 1984) and has been documented in many African and Asian elephant populations (Dublin, 1983; Eisenberg, 1980; Gadgil & Nair, 1984; Jayantha et al., 2009; Lee, 1987; Lee & Moss, 2011; Rapaport & Haight, 1987; Vidya, 2014; Woodford & Trevor, 1970). However, there are only a few detailed studies of calf-conspecific interactions in elephants (Lee, 1987; Lee & Moss, 2011 on African savannah elephants (*Loxodonta africana*); Gadgil & Nair, 1984 on semi-captive Asian elephants (*Elephas maximus*); Chelluri 2009 on wild African forest elephants (*Loxodonta cyclotis*); Webber 2017 on wild Asian elephants in Uda Walawe, Sri Lanka, and captive Asian and African savannah elephants) and hardly any studies quantifying allomaternal care and detailing how such care results in calves experiencing differentiated relationships (but see Lee, 1987; Lee & Moss, 2011).

We, therefore, investigated the ontogeny of social relationships with a focus on caretaking behaviours in a long-term monitored wild Asian elephant population in southern India (Kabini Elephant Project: Keerthipriya and Vidya, 2022; Vidya et al., 2014). We did so by comparing behaviours of different categories of subadult and adult females towards calves up to 6 months of age and *vice versa*. Specifically, we examined 1) whether proximity contacts and interactions were initiated and terminated primarily by adult and subadult female conspecifics or calves (a, d, and f in Results), 2) how care (and hence, calf-conspecific interactions) was distributed among female conspecifics in the group and whether individuals that showed one form of care (escorting, see below) also provided other forms of care (b, c, d, f in Results), and 3) whether certain kinds of behavioural interactions were associated with specific categories of conspecifics (e in Results).

In our study, we focused on calves <6 months old, as significant behavioural development commences during the first six months of life (Revathe et al., 2020). This critical developmental window is also marked by high mortality (Kabini Elephant Project, unpublished data), necessitating heightened care. The latter might be expected to result in female conspecifics initiating proximity contacts and interactions, whereas the precociality hypothesis (Hill & Campbell, 2014) predicted that calves would initiate such contacts and interactions as they are precocial and quick to explore their surroundings after birth (Nair, 1989; Revathe et al., 2020). It was previously found in the Kabini population that certain non-mother females showed coordinated movement with particular calves, beyond that resulting from coordinated group movement (Revathe, 2022). We termed these females ‘escorts’. In this population, at least one escort was present in around three-fourths of the female group sightings that had a calf (Revathe, 2022). Because of differentiated relationships among adult females (Nandini et al., 2017, 2018) and the consistent presence of specific escorts for specific calves across elephant groups (Revathe, 2022), we predicted that calves would have differentiated social relationships with the conspecific females in their clan from a young age. We expected calves to initiate more spatial and behavioural interactions with their mother – the reliable/familiar social partner, source of nourishment for the calf, and female with the highest incentive to show positive interactions to the calf – than with other conspecific categories of females in the group, and *vice versa*. We did not have an *a priori* expectation about whether escorts would show more affiliative behaviour towards calves, and *vice versa*, than would other females in the group. As being near calves by coordinating movement might be thought of as a form of care by itself, it was not necessary that escorts should show other forms of care, and other individuals could potentially provide such care. We did not expect the trajectory of social ontogeny to diverge between females and males within the first six months of life based on the similar pattern of development across sexes in calves (up to 1 year old) (Revathe et al., 2020). However, there could be an effect of age on calf relationship differentiation, as calves start to explore their physical and social environment.

In essence, to gain a richer understanding of social ontogeny, we examined certain behaviours (initiation and termination of proximity and interactions and responses to interactions) from the perspective of both the calves and conspecific females, as it takes two to form a relationship. Such an approach can help clarify the relative contributions of individuals (i.e., calf vs conspecific female vs both; see Hinde & Spencer-Booth, 1967) to the development of differentiated social relationships.

## Methods

### Field data collection

We collected field data from December 2015 to June 2018 in Nagarahole and Bandipur National Parks and Tiger Reserves, southern India, where the tiger density is high (Jhala et al., 2008). We carried out field sampling across these two parks following systematic, fixed sampling routes, from ∼6:30 AM to ∼5:45-6:45 PM (see Nandini et al., 2017 for further details). Female elephant groups were defined as one or more females, often accompanied by young ones, that showed coordinated movement or behaviour and were usually within 50-100 m of one another (Nandini et al., 2017, 2018; equivalent to ‘party’ in the primate literature). We aged, sexed, and identified the individuals seen based on natural physical characteristics (Vidya et al., 2014). The Kabini Elephant Project has recorded hundreds of individually identified elephants since its inception in 2009 (see Vidya et al., 2014). We age-classified females as subadults (5-<10 years) and adults (>=10 years). Since we were interested in the development of social relationships, we examined calves of different ages. We categorized calves according to differences in behavioural development based on a previous study on this population (Revathe et al., 2020). Therefore, in this study, we refer to those less than three months old as newborn calves and to those between 3 and <6 months of age as infant calves.

The subjects of the study were 20 calves ≤6 months old, belonging to 9 clans. Most of the behavioural data presented here were collected from around the Kabini backwaters (which separate Nagarahole and Bandipur) because of good visibility. Thus, we sampled clans that used this area. We used focal animal sampling (Altmann, 1974) to record interactions between calves and conspecifics within groups. We recorded observations using a video camera (SONY HDR-XR 100E or SONY HDR-PJ 540E).

We defined escorts based on coordinated movement between calf-conspecific pairs, beyond that required of group members (i.e., any distance less than 50-100 m that the group showed) (Revathe, 2022). Thus, one or more non-mother subadult or adult females who moved along with the focal calf (either accompanied by the calf’s mother or not) was/were identified as escort(s) in each focal session (Figure 1a) as previously done in whales (e.g., Gero et al., 2009). We refer to females other than the focal calf’s mother and escort(s) in a focal session as ‘other females’ (Figure 1a). Based on this qualitative difference observed in the field (Revathe, 2022), we differentiated between escorts and other females because not acknowledging this distinction and clubbing all conspecific females, other than the mother, into one category could mask the presence of differentiated social relationships, if present, of calves with conspecific females.

**Figure 1.**
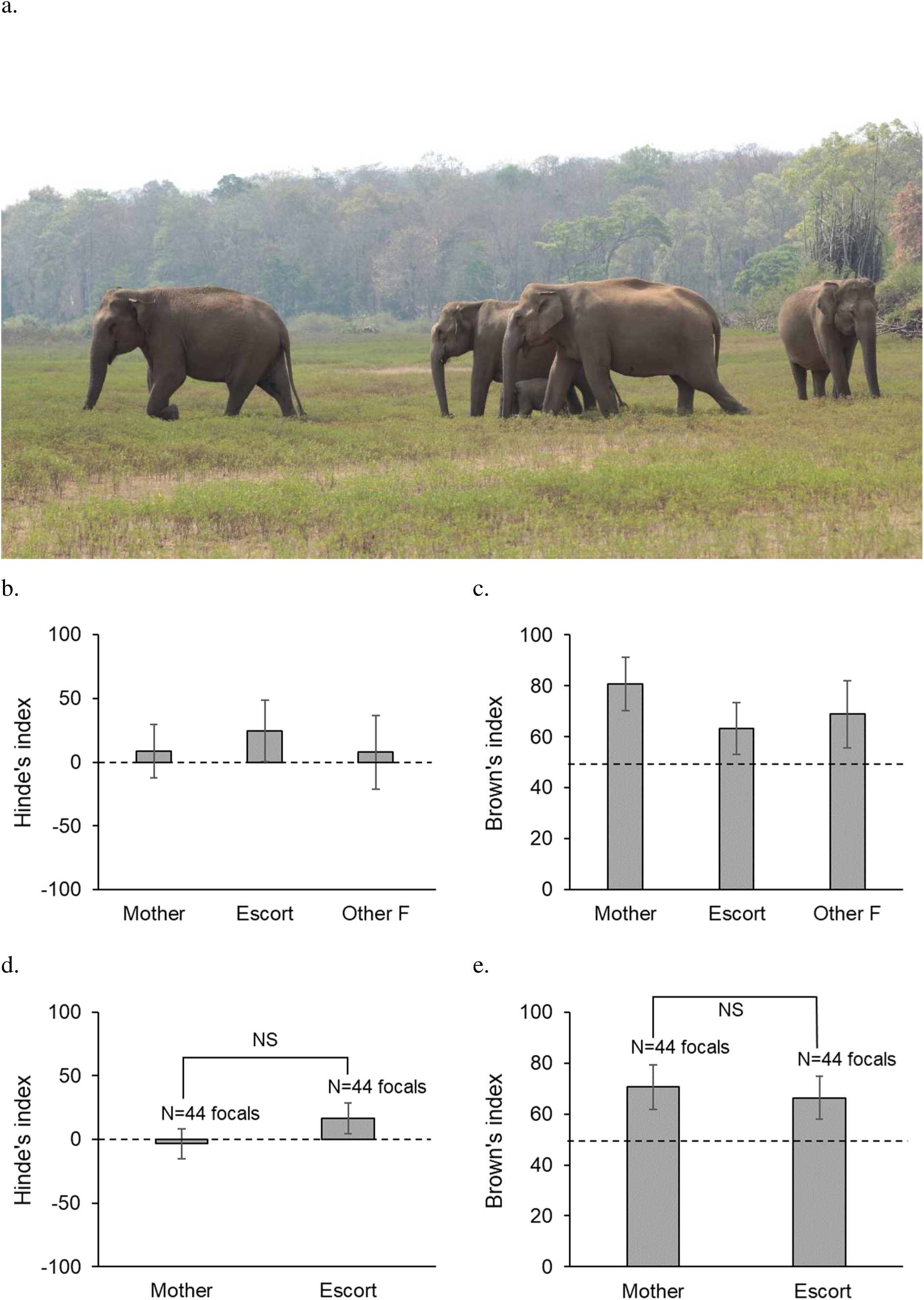
a) Photo of an elephant group showing the positions of group members. Left to Right: other female, mother, calf, escort, other female. Photo: Divya Choudhary, © Kabini Elephant Project; b) average Hinde’s and c) average Brown’s proximity indices of 7 calves for which data were available with all three conspecific categories; d) Hinde’s and e) Brown’s indices of 17 calves with mothers and escorts. Error bars are 95% CI. Dashed lines indicate the expected Hinde’s index of 0 and expected Brown’s index of 50 in the respective graphs.

Although an escort is a female that showed coordinated movement with the calf, she did not have to necessarily initiate any, and specifically, allocare, interaction towards the calf. Mother-escort comparisons in their behaviours towards calves and *vice versa* allowed us to understand how comparable allomaternal care (if shown by escorts) was to maternal care, and if it was primarily the escorts, and not other females, that showed allomaternal care (we defined this to require more than coordinated movement with the calf). It should be noted that although, by definition, other females did not show coordinated movement with the focal calf, they were still part of the same group as the focal calf and, hence, could reach the calf to initiate (allocare) interactions (Figure 1a). Similarly, it was possible for calves to easily go up to other females to initiate interactions. Essentially, it was entirely feasible for conspecific females to show escorting behaviour but not allomaternal care and *vice versa*. Data could not be recorded blindly because our study involved focal animals in the field.

Fieldwork was carried out with relevant Forest Department permissions and in accordance with India’s wildlife protection laws.

### Focal video scoring and data analysis

We used Windows Media Player 12.0 to score focal videos to quantify calf-conspecific proximity and interactions (see below). All the videos were scored by TR. We used only those focals in which all the group members were clearly visible, and in which all three conspecific categories of females (Mother, Escort, Other female) were present for the focal calf because we wanted to make mother-escort, mother-other female, and escort-other female comparisons. Since a majority of the calves born during the study period had at least one escort in almost all their sightings (Revathe, 2022) and as calves without an escort often did not survive past a few months (Kabini Elephant Project, unpublished data), it was not feasible to obtain multiple focals of calves without an escort. However, since such calves are only a minority in the study population, we do not expect the lack of their focals in the analyses presented below to substantially affect our results. Importantly, we use our results to only infer whether calves with an escort develop differentiated relationships and whether escorting behaviour could be used as a proxy to identify allomothers in the field. It is likely that calves without an escort also eventually have differentiated social relationships, however that is beyond the scope of the current study.

### Calf-conspecific proximity contacts

We scored 3 focal videos per calf, recorded on three different days on a second-by-second basis for 20 calves who were up to 6 months of age, and noted down who initiated movements (approach or leave; *N*=299 movements) (Supplementary Material 1). The calf belonged to the same age-class during all 3 focals, and each calf appeared in only one age-class. Each focal lasted 20 minutes (72,000 seconds scored in all), during which we noted down the focal calf’s positions in terms of who was near the calf (see Supplementary Material 2). We considered a focal calf to be near a conspecific if the distance between the calf and the conspecific was up to one calf-body length (≤1 m). A calf could be near multiple individuals simultaneously.

We used the Hinde’s proximity index (Hinde & Spencer-Booth, 1967; the percentage of approaches made by the calf towards a conspecific minus the percentage of leavings made by that calf away from that conspecific) to find out whether the calf more frequently approached or left a particular conspecific (see Supplementary Material 3). Additionally, we calculated Brown’s proximity index (Brown, 2001) to measure the relative contributions of the calf and conspecific to changes in proximity contacts between the pair (see Supplementary Material 3).

Since these indices could be calculated with all three conspecific categories in each focal only for two calves (as the remaining did not show proximity contact changes with other females), we could not use parametric tests. We instead tested for differences in Hinde’s and Brown’s indices across conspecific categories (Mother, Escort, Other Female) using the non-parametric Friedman’s ANOVA, using the averaged value across focals for each calf and matched for calf identity. We additionally performed ANOVAs on the logit Hinde’s index and Brown’s index to examine possible differences between only mothers and escorts, for which the sample sizes were larger. Here, conspecific category was a fixed factor with two levels – Mother and Escort. Calf identity was included as a random factor because there were multiple calves, although each calf was sampled only in two or three focals. However, we were not actually interested in variation amongst calves, nor can such variation be tested without many focals per calf. Therefore, in this and all the other following analyses whenever calf identity was used, inferences were not made about calf identity or its interactions. Only calves that had Hinde’s and Brown’s index values in at least two of their focals were used. Due to low sample size, the effects of calf age and sex on proximity initiation could not be assessed.

### Calf-conspecific proximity

During each 20-minute focal, we took a scan every 4 minutes to obtain six independent calf positions in terms of the conspecific(s) who were near the calf (4 minutes was chosen based on how quickly calves changed their positions: this corresponded to 95% of the durations to changes in positions; see Supplementary Material 2). We thus used a total of 360 calf positions (6 positions x 3 focals x 20 calves, of which 10 calves were newborns, less than 3 months old, and 10 calves were infant calves, 3-<6 months old) for this analysis. In order to find out whether calf proximity differed with respect to conspecific category, we carried out an ANOVA on the logit proportion of calf-scans near a conspecific (dependent variable; proportions were logit transformed as they were not normally distributed), with calf age-class (<3 and 3-<6 months) and conspecific category (Mother, Escort, and Other Female) as fixed factors, and calf identity nested within age-class as a random factor.

### Calf-conspecific interactions

To examine calf-conspecific interactions, we scored two 30-minute focal videos per calf for the same 20 calves as those used for proximity analyses, during which we noted down all the interactions that occurred between a focal calf and females of the three conspecific categories, the identities of the initiator and terminator of each interaction, and the duration of the interaction. Behaviours were classified into three classes: feeding-related (sucking from a conspecific, taking plucked grass from a conspecific, sniffing grass from the mouth of a conspecific, etc.), resting-related (calf leaning against a conspecific, calf lying down near/under a conspecific, and calf sliding off an individual to lie down), and social (for e.g., touching a conspecific, rubbing against a conspecific, conspecific standing guard over a calf etc.; see Supplementary Material 4). We also noted down whether conspecific-initiated interactions, as well as the responses shown by conspecifics towards calf-initiated interactions, were positive (for e.g., rushing towards a calf when the calf was in distress, stopping one’s activity to allow the calf to suck, keeping the calf within reachable distance, etc.), negative (for e.g., lashing out at a calf, kicking a calf, pushing a calf, etc.), or neutral (see Supplementary Material 4). Conspecifics did not always show a response, and ‘no response’ was also included under the category ‘neutral’ during analysis.

#### Initiation of interactions

In order to find out whether calves initiated more interactions towards conspecifics or *vice versa* and to find out whether there was a difference amongst mothers, escorts, and other females in the numbers of interactions that calves initiated towards them and the numbers of interactions that they initiated towards calves, we ran a nested ANOVA on the log-transformed numbers of calf-conspecific interactions (as they were not normally distributed), with calf age-class (<3, 3-<6 months), initiator category (Calf, Conspecific), and conspecific category (Mother, Escort, and Other Female) as fixed factors, and calf identity nested within age-class as a random factor. We also tested interaction effects.

#### Conspecific-initiated interactions: frequency in different contexts

We then classified conspecific-initiated interactions towards calves into four different interaction types (Supplementary Material 4, Table 2) from the perspective of the calf, namely positive (potentially benefitting the calf; such as checking the calf, chaperoning the calf), neutral (no obvious benefit to the calf; such as sniffing the calf), and negative (not benefitting the calf, or at least not in the immediate interest of the calf). Since not all the negative interactions cause physical distress, we also split them into non-aggressive negative (not causing physical distress, such as nudging the calf to move, mother nudging the calf towards the escort) and aggressive negative (causing physical distress, such as kicking the calf, lashing out at the calf) interactions. We found that most of the calf-conspecific interactions were initiated by calves (see Results); hence, we could not use ANOVA to test the effect of conspecific category on interaction type with calves. We, therefore, used an R x C test of independence using *G*-test to test the null hypothesis that interaction type with calves was independent of conspecific category by combining interactions across all focals (of all calves) in each conspecific category and interaction type.

#### Calf-initiated interactions: frequency in different contexts, and terminations and types of responses shown by conspecifics

To understand social preferences of calves, we examined whether the number of calf-initiated interactions varied based on conspecific category for the different behavioural classes. To do so, we carried out a nested ANOVA on the log-transformed numbers of calf-initiated interactions, with calf age-class, conspecific category, and behavioural class of interaction (Feeding, Resting, and Social) as fixed factors, and calf identity nested within age-class as a random factor.

For each focal, we also calculated the proportion of calf-initiated interactions towards females of a conspecific category terminated by females of that category (calves that did not initiate any interaction towards females in a focal were not included in the analysis, *N*=2 calves in each age-class). These proportions of calf-initiated interactions terminated by mothers and escorts were compared. Terminations by other females were not used as only two calves initiated interactions with all three conspecific categories of females in all their focals. We carried out a nested ANOVA on the logit-transformed proportions of calf-initiated interactions terminated by conspecifics, with calf age-class (<3 and 3-<6 months) and conspecific category (Mother, Escort) as fixed factors and calf identity nested within age-class as a random factor.

We also calculated the proportions of calf-initiated interactions towards mothers and escorts that elicited positive, neutral, or negative responses from that conspecific category (these proportions would add up to 1 for each conspecific category; therefore, the proportion of negative responses was not included in the analysis). Again, as above, responses by other females could not be compared as calves did not initiate sufficient interactions towards them. To see if mothers and escorts showed similar types of responses, we ran a nested ANOVA on the logit proportions of calf-initiated interactions that elicited a positive or a neutral response, with calf age-class, conspecific category, and type of response (Positive, Neutral) as fixed factors, and calf identity nested within age-class as a random factor.

Thus, to check whether females of different conspecific categories behaved in a similar manner towards calves, we compared conspecific-initiated interactions towards calves, proportion of calf-initiated interactions that were terminated by conspecifics, and the proportion of calf-initiated interactions that elicited different kinds of responses, across the conspecific categories. To check whether calves behaved in a similar manner towards different categories of conspecific females, we compared calf proximity initiation and leaving, calf proximity, and the number of calf-initiated interactions – overall and under different behavioural classes, across the conspecific categories.

### Calf-conspecific non-suckling interactions

It was possible that nutritional dependence of calves on their mother might mask the presence of otherwise comparable relationships of calves with escorts or other females. To investigate this possibility, we modified the original dataset by removing all the suckling-related interactions that the focal calves initiated towards the conspecifics and investigated whether the pattern of social interactions of calves remained the same even when suckling was not considered. The same analyses that were performed on calf-initiated interactions were also performed on this modified dataset. We report these results in the supplementary materials and present only graphs and *P*-values in the results in the main text.

The ANOVAs were carried out by obtaining sums of squares using Statistica (7.0, StatSoft, Inc. 2004), and the *F* test calculations were carried out based on Neter *et al*. (1990, Chapter 27, pgs. 1010-1029).

## Results

### a) Initiatiation and termination of proximity contacts by calves and conspecifics, and differences across conspecific categories (mothers, escorts, other females) in proximity contact initiation

There was no significant difference in the Hinde’s (Friedman ANOVA: χ^2^= 2.000, *N*=7 calves, *df*=2, *P*=0.368, Figure 1b) or Brown’s (Friedman ANOVA: χ^2^= 4.571, *N*=7 calves, *df*=2, *P*=0.102, Figure 1c) proximity indices among mothers, escorts, and other females. ANOVAs on the logit Hinde’s index with mothers and escorts based on data from a larger number of calves (17 calves, 44 focals) also showed no significant effect of conspecific category (*F*_1,16_=3.849, *P*=0.067, Figure 1d). However, the average Hinde’s index of the calves with their mothers was not significantly different from zero, suggesting that calves approached and left their mothers to similar extents, while the Hinde’s index with the escorts was significantly greater than zero, suggesting that calves were more likely to approach than to leave escorts (confidence intervals in Figure 1d). The average Brown’s proximity indices of calves with the mother, escort, and other females were greater than 50 (Figure 1c,e), indicating that calves contributed more than their conspecifics to changes in proximity. However, there was no significant effect of conspecific category (mothers and escorts) on this index either (ANOVA on logit Brown’s index: *F*_1,16_=0.888, *P*=0.360; 17 calves, 44 focals; Figure 1d).

### b) Effects of calf age-class and conspecific category on calf-conspecific proximity

Calves were within easy reachable distance (≤1 m) of at least one female conspecific almost all the time (average proportion of scans near at least one female conspecific ± 95% CI: newborn calves: 0.978 ± 0.022; infant calves: 0.930 ± 0.030). There was no significant effect of calf age-class on the proportion of scans when calves were near conspecific females (ANOVA: *F*_1,18_=0.205, *P*=0.656), with newborn calves and infant calves spending similar proportions of scans near conspecific females (average of the proportion of scans near mother, escort, and other females ± 95% CI = 0.475 ± 0.076 for newborn calves and 0.439 ± 0.069 for infant calves; these values were lower than the average proportion of scans near at least one female conspecific because of averaging across conspecific categories, with other females not spending much time near calves). However, there was a significant main effect of conspecific category (*F*_2,36_=103.315, *P*<0.001), and its interaction with calf age-class (*F*_2,36_=8.139, *P*=0.001; Figure 2).

**Figure 2.**
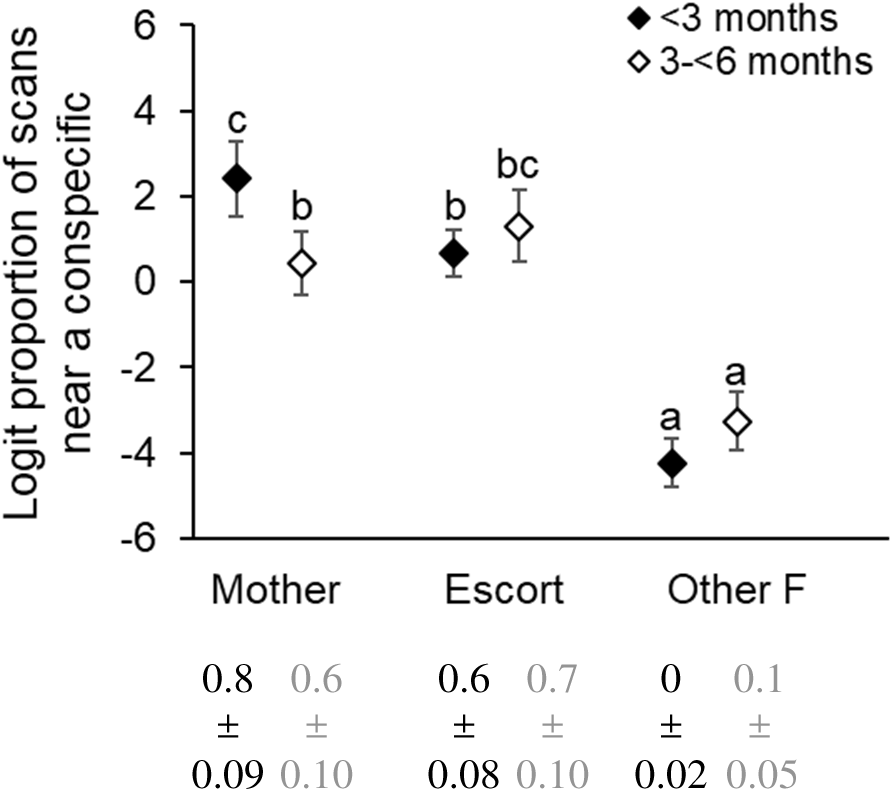
Logit proportions of scans that a calf spent near the three conspecific categories of females for 20 calves (<3 months: *N*=10 calves; 3-<6 months: *N*=10 calves). Error bars are 95% CI. In this and the following graphs, letters above the data points indicate pattern of statistical significance based on Tukey’s HSD tests (a<b<c), unless otherwise specified. Shared letters above the bars indicate no significant difference between comparisons. The untransformed value (average ± 95% CI) is shown below the graph for each category. (Averaging was done over all the focals of all the calves in the category in this and the following graphs.)

The proportions of scans in which calves were near their mothers or escorts (which were not significantly different from each other; *P*=0.529) were significantly higher than that near other females (mother versus other females: *P*<0.001; escort versus other females: *P*<0.001). Newborn calves spent a greater proportion of their time near mothers than escorts (*P*=0.043) or other females (*P*<0.001), and a greater proportion of their time near escorts than other females (*P*<0.001; Figure 2). Infant calves spent a similar proportion of their time near mothers and escorts (*P*=0.642), but these were higher than that spent near other females (*P*<0.001 for both the comparisons, Figure 2). Newborn calves spent a significantly greater proportion of time near their mothers than infant calves did (*P*=0.015), but both spent similar proportions of time near their escorts (*P*=0.871) and near other females (*P*=0.529; Figure 2). When tested for infant calves separately, there was no effect of calf sex on the proportion of scans spent near conspecifics (Supplementary Material 5).

### c) Differences across conspecific categories in the number and nature of interactions (i.e., positive / neutral / negative) initiated towards calves

We observed 93 positive, 10 neutral, 35 non-aggressive negative, and 10 aggressive negative conspecific-initiated interactions towards calves. The non-aggressive negative interactions included only nudge (to make the calf move from a spot), whereas the aggressive negative interactions included kick, push, pull trunk, beat with tail, and lash (Supplementary Material 4, Table 1). We found that the type of conspecific-initiated interactions towards calves was dependent on conspecific category (*G*_corrected_ = 45.930, *P*<0.001). Escorts initiated a majority (77.4%) of the 93 positive interactions towards calves, mothers initiated 20.4% of them, and other females initiated only 2.2% of them. Escorts also initiated most of the 10 neutral interactions (80%) towards calves, whereas mothers and other females initiated only one neutral interaction each in total. Escorts also initiated a majority (60%) of the 35 negative non-aggressive interactions towards calves, followed by mothers (31.4%), and other females (8.6%). Aggressive negative interactions towards calves were rare (10 interactions), and mostly (90%) initiated by other females; one interaction was initiated by a mother and none by an escort.

**Table 1.**
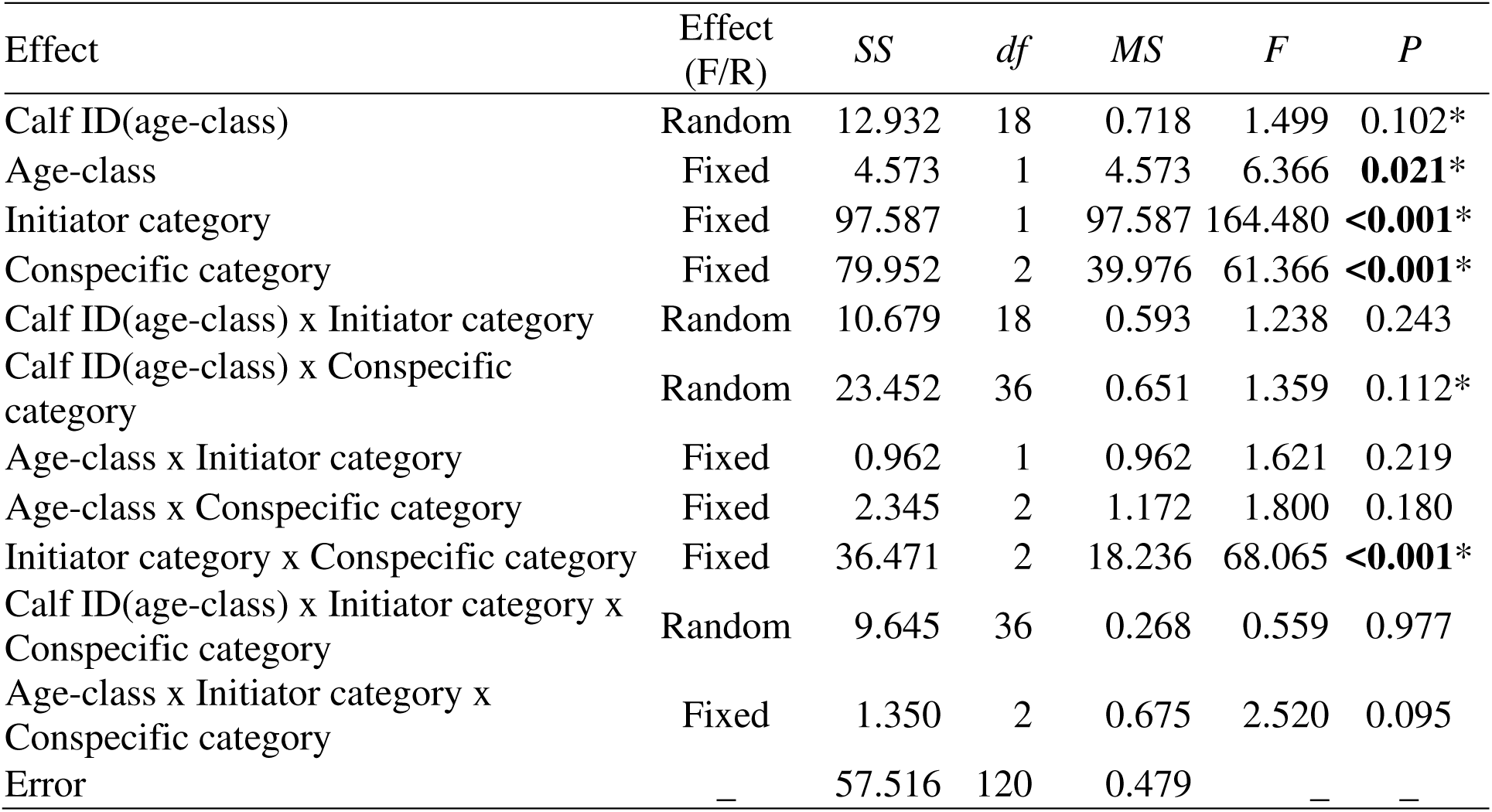
Results of the nested ANOVA on the log numbers of all calf-conspecific interactions. Calf ID (random factor) was nested within age-class, and age-class, initiator category, and conspecific category were fixed factors. Significant *P* values are marked in bold. The asterisks in the *P* values column indicate significance in the ANOVA on the log numbers of non-suckling calf-conspecific interactions (see Supplementary Material 8) for comparison. (Inferences should not be drawn about Calf ID or its interactions in this and the following tables as each calf was sampled in only two focals.)

### d) Initiation of behavioural interactions by calves and conspecifics

Of the total of 1332 interactions between calves and their mothers, escorts, and other females in their groups that we observed during the total focal duration of 1200 minutes, calves initiated 1184 interactions. Most of these interactions were of very short duration (average duration of interaction ± 95% CI: calf-initiated: 20.4 ± 4.14 seconds; conspecific-initiated: 24.0 ± 14.5 seconds; Supplementary Material 6). We found significant main effects of initiator category, calf age-class, and conspecific category on the log number of calf-conspecific interactions (Table 1).

Calves initiated significantly more interactions towards conspecific females (average number of interactions ± 95% CI per 30-min focal: 29.60 ± 4.931) than conspecific females initiated towards calves (3.70 ± 1.634; Table 1, Figure 3a). Newborn calves were involved in a significantly greater number of interactions (39.4 ± 8.87) with conspecific females (initiated by either) than were infant calves (27.3 ± 6.64; Table 1). The numbers of calf-mother (14.88 ± 3.644) and calf-escort (16.95 ± 3.814) interactions were similar (95% CI around difference between means for conspecific category: 0.312; Tukey’s HSD: *P*>0.05), and both were significantly higher (*P*<0.05 for both) than the number of calf-other female interactions (1.48 ± 0.848).

**Figure 3.**
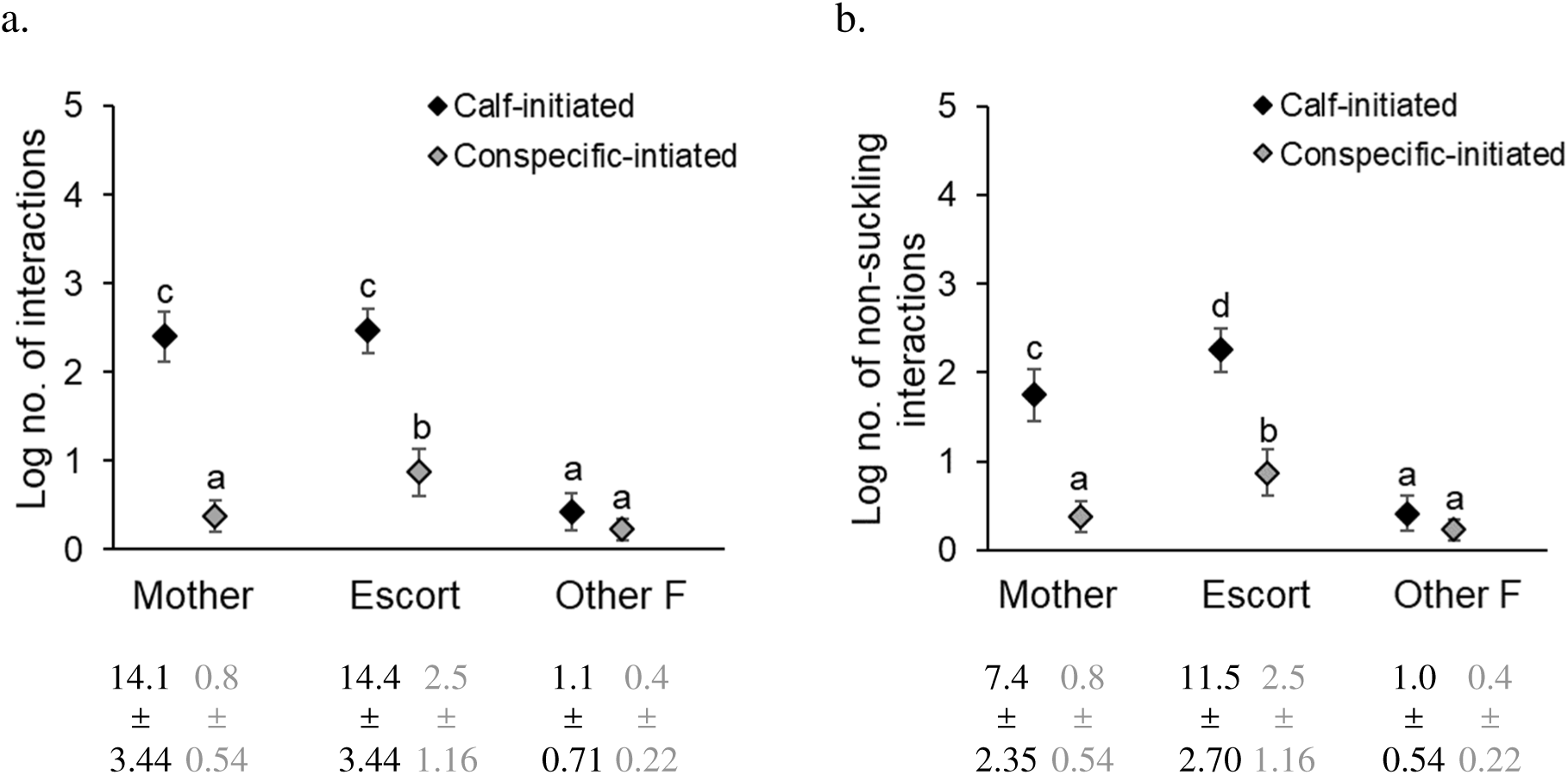
Log numbers of a) all interactions and b) non-suckling interactions initiated by calves towards the three conspecific categories of females, and the log numbers of interactions initiated by these conspecific females towards calves per focal for 20 calves (<3 months: *N*=10 calves; 3-<6 months: *N*=10 calves). Error bars are 95% CI. The untransformed number of interactions (average ± 95% CI) for each category is shown below the X-axis.

There was also a significant interaction effect between initiator and conspecific categories (Table 1). Calves initiated similar numbers of interactions towards mothers and escorts (95% CI around difference between means for initiator category x conspecific category: 0.348; *P*>0.05; Figure 3a), and a smaller number of interactions towards other females (*P*<0.05 for both the comparisons; Figure 3a). On the other hand, escorts initiated a greater number of interactions towards calves than mothers or other females did (*P*<0.05 for both the comparisons; previous paragraph, Figure 3a), with the latter two not being significantly different from each other (*P*>0.05; Figure 3a). When tested separately, there was no effect of calf sex on the numbers of calf-conspecific interactions involving infant calves (Supplementary Material 7).

### e) Calf initiation of interactions of certain behavioural classes (feeding / resting / social) towards conspecific categories

Among the 1184 calf-initiated interactions were 539 feeding interactions, 89 resting interactions, and 556 social interactions. The log number of calf-initiated interactions was significantly affected by the behavioural class of interaction, conspecific category, their interaction, and calf identity (Table 2). Calves initiated more interactions towards mothers and escorts than towards other females as seen above (Initiator category x Conspecific category in Table 1, Figure 3a). This pattern was also found separately in feeding (95% CI around difference between means for behavioural class x conspecific category: 0.502; Mother versus Other Females: *P*<0.05; Escort versus Other Females *P*<0.05, Figure 4a) and social interactions (Mother versus Other Females: *P*<0.05; Escort versus Other Females *P*<0.05, Figure 4a), but the number of resting-related interactions initiated by calves was higher only towards escorts than towards other females (*P*<0.05), and was similar between mothers and other females (*P*>0.05) and mothers and escorts (*P*>0.05, Figure 4a).

**Table 2.** Results of the nested ANOVA on the log number of calf-initiated interactions towards the three conspecific categories of females. Significant *P* values are marked in bold. The asterisks in the *P* values column indicate significance in the ANOVA on the log numbers of calf-initiated non-suckling interactions (see Supplementary Material 11) for comparison.

| Effect | Effect (F/R) | *SS* | *df* | *MS* | *F* | *P* |
| --- | --- | --- | --- | --- | --- | --- |
| Age-class | Fixed | 1.151 | 1 | 1.151 | 1.153 | 0.297 |
| Calf ID(Age-class) | Random | 17.973 | 18 | 0.999 | 2.171 | **0.005\*** |
| Behavioural-class | Fixed | 52.362 | 2 | 26.181 | 27.138 | **<0.001\*** |
| Conspecific category | Fixed | 86.172 | 2 | 43.086 | 59.997 | **<0.001\*** |
| Behavioural-class* Conspecific category | Fixed | 22.078 | 4 | 5.519 | 11.206 | **<0.001\*** |
| Age-class*Behavioural-class | Fixed | 1.379 | 2 | 0.689 | 0.715 | 0.496 |
| Age-class*Conspecific category | Fixed | 0.539 | 2 | 0.269 | 0.375 | 0.690 |
| Age-class*Behavioural-class*Conspecific category | Fixed | 2.166 | 4 | 0.542 | 1.100 | 0.363 |
| Calf ID(Age-class)*Behavioural-class | Random | 34.730 | 36 | 0.965 | 2.098 | **0.001\*** |
| Calf ID(Age-class)*Conspecific category | Random | 25.853 | 36 | 0.718 | 1.561 | **0.031\*** |
| Calf ID(Age-class)*Behavioural-class*Conspecific category | Random | 35.464 | 72 | 0.493 | 1.071 | 0.353 |
| Error | — | 82.788 | 180 | 0.460 | — | — |

**Figure 4.**
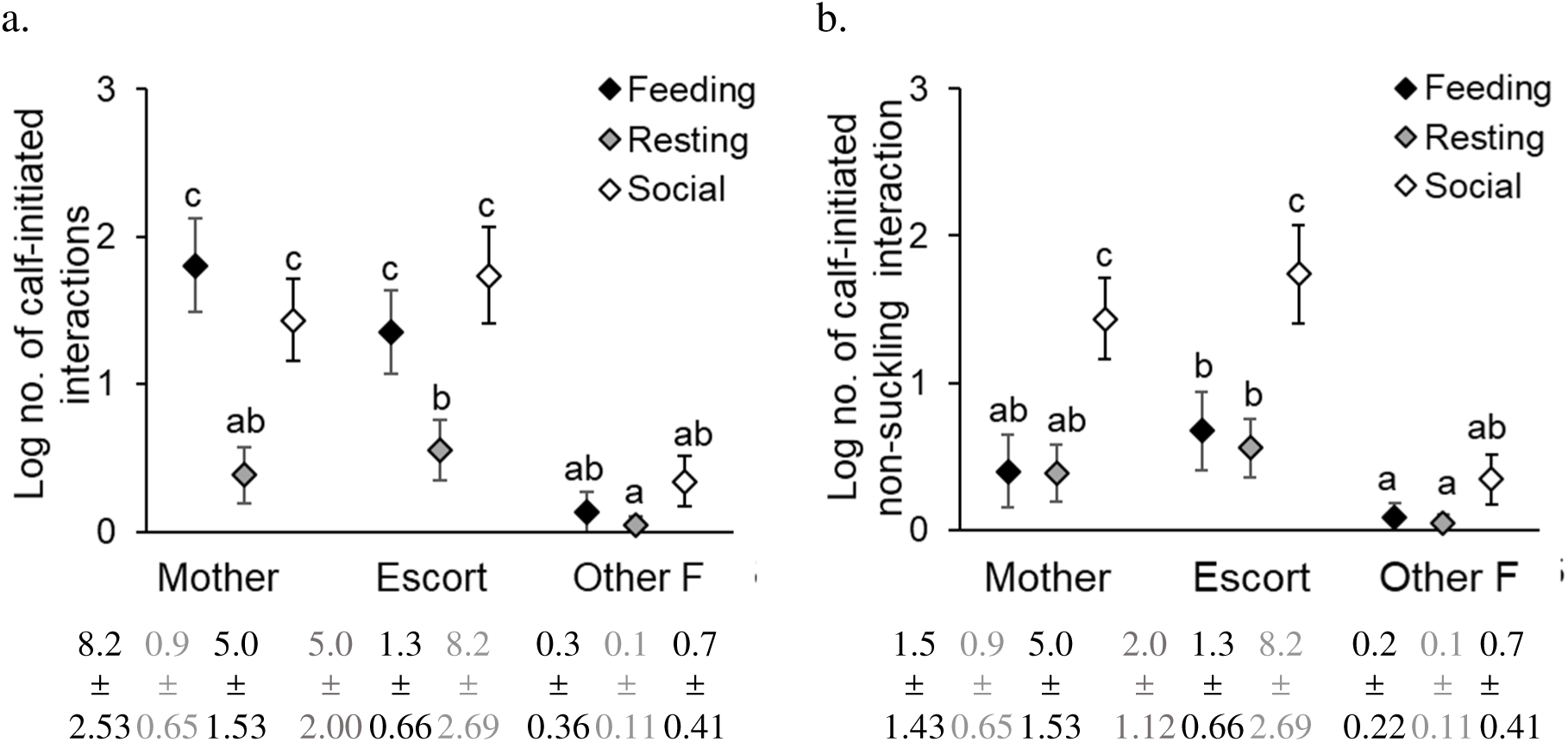
Log numbers of a) all calf-initiated interactions and b) calf-initiated non-suckling interactions of three behavioural classes towards the three conspecific categories of females for 20 calves (<3 months: *N*=10 calves; 3-<6 months: *N*=10 calves). Error bars are 95% CI. Untransformed numbers (average ± 95% CI) are shown below the graphs.

Newborn and infant calves initiated similar numbers of interactions towards conspecific females (in keeping with the lack of an Age-class x Initiator category effect in Table 1). Calves initiated similar numbers (average ± 95% CI per 30-minute focal) of feeding (13.48 ± 3.473) and social interactions (13.90 ± 3.308; 95% CI around difference between means for behavioural class: 0.310; Tukey’s HSD: *P*>0.05) towards conspecific females, which were both significantly higher than resting-related interactions (2.23 ± 1.208; *P*<0.05 for both the comparisons); this pattern was not different across calf age-classes (Table 2). There was no effect of sex on the number of calf-initiated interactions for infant calves examined separately (see Supplementary Material 10).

### f) Termination of interactions by calves and conspecifics, and differences across conspecific categories in the kinds of responses shown towards calf-initiated interactions

#### Terminations

Calves themselves terminated most (*N*=868) of the interactions that they initiated (*N*=1184), while conspecifics terminated the remaining. We found that mothers and escorts terminated similar proportions of calf-initiated interactions towards them (*P*=0.271; Figure 5, Supplementary Material 12). The same pattern was seen in the termination of non-suckling interactions (Supplementary Material 12). In both the cases, mothers and escorts terminated similar proportions of interactions initiated by newborn and infant calves.

**Figure 5.**
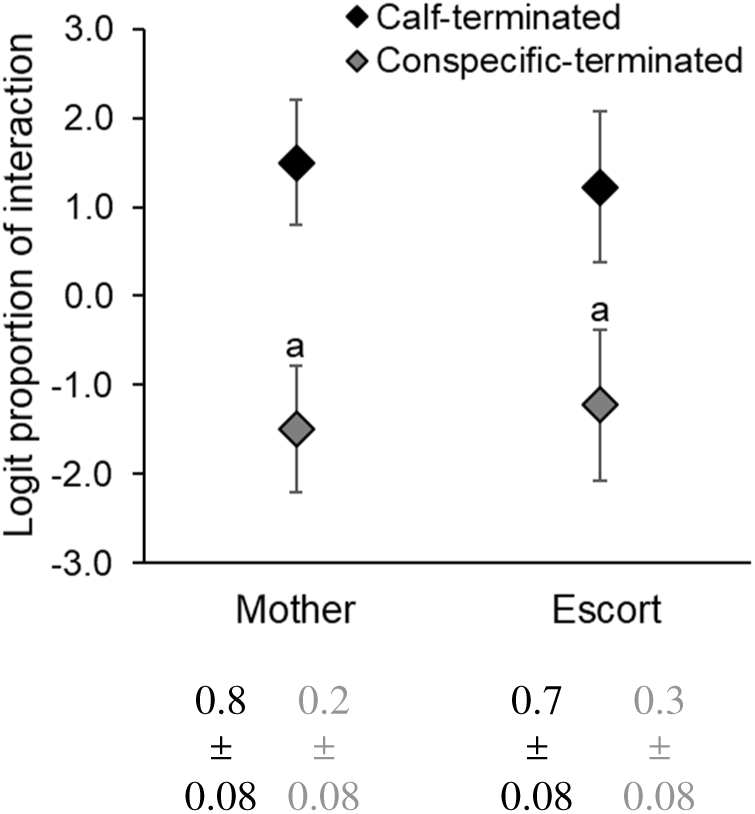
Logit proportions of calf-initiated interactions with their mothers and escorts that were terminated by the calves and by the conspecific females for all calf-initiated interactions for 16 calves (<3 months: *N*=8 calves; 3-<6 months: *N*=8 calves). Error bars are 95% CI. Letters indicate pattern of statistical significance based on ANOVA. Proportions of calf-initiated interactions terminated by the calf and mother, and by the calf and escort, respectively, (average ± 95% CI) are shown below the graph.

#### Responses by conspecifics

Calves received different types of responses, but most of their interactions did not elicit a response from conspecific females (885 out of 1184 calf-initiated interactions received no response). There were more negative (192) than positive (107) responses overall, but only 17 of the 192 negative responses were aggressive, such as kick, lash, push, pull trunk, and beat with tail.

We found that the proportions of calf-initiated interactions towards mothers and escorts that elicited a neutral response (about 80%) were significantly higher than the proportion that elicited a positive response from them (about 10%; *P*<0.001; Figure 6, Supplementary Material 13). There was no significant effect of conspecific category or its interaction with response type (Figure 6, Supplementary Material 13) on the proportion of calf-initiated interactions that elicited a response.

**Figure 6.**
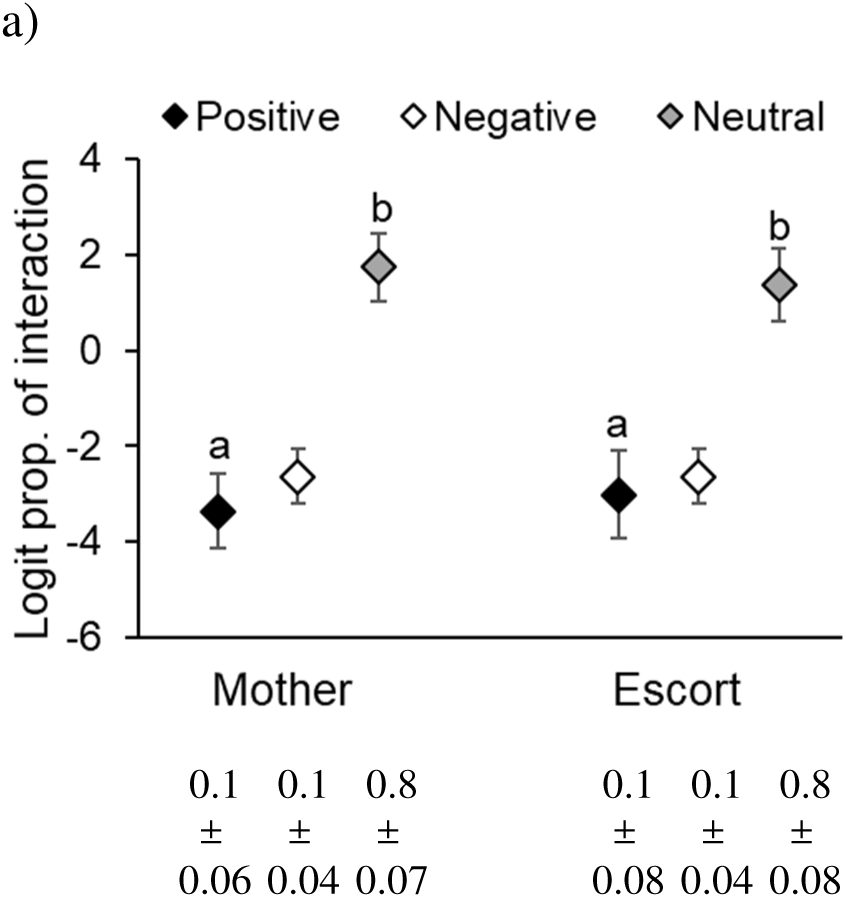
Logit proportions of a) all calf-initiated interactions and b) non-suckling calf-initiated interactions with their mothers and escorts that elicited a positive, neutral, and negative response from them for 16 calves (<3 months: *N*=8 calves; 3-<6 months: *N*=8 calves). Error bars are 95% CI. Letters above the data points indicate patterns of statistical significance (a<b). Proportions of interactions that elicited a positive, neutral, and negative response from mothers and escorts are written below the graph (average ± 95% CI).

When we pooled together negative responses to calf-initiated interactions and negative conspecific female-initiated interactions towards calves across focals, we found that calves less than 6 months of age received the lowest rate of aggression from their mothers (0.15/h), followed by escorts (0.4/h). Calves received the highest rate of aggression from other females (0.8/h), despite calves spending the least proportion of time near other females.

## Discussion

We examined the relationship between calves that were up to 6 months old and conspecific females (that were at least 5 years old) to understand the ontogeny of social relationships in wild Asian elephants. Through analysing calf-female spatial and behavioural interactions, we found that calves were precocial in forming new relationships. Differentiated social relationships emerged through escorting females also exhibiting other positive behavioural interactions with the calf as much as or more than the calf’s mother, through mothers and escorts being similar in their behaviours towards calves and calves towards them, and through the presence of other females who engaged in more negative than positive interactions with calves. Thus, our study on social ontogeny shows that differentiated social relationships are formed from a young age in Asian elephants.

### Elephant calves are precocial in the formation of new social relationships

#### Proximity

We found that calves were responsible for initiating more changes in proximity with mothers, escorts, and other females than *vice versa* from a young age, in line with the predictions of the precociality hypothesis (Hill & Campbell, 2014). Offspring have been found to contribute more to proximity changes than mothers in other precocial species such as belugas (*Delphinapterus leucas*; Hill & Campbell, 2014), Saharan arrui (*Ammotragus leruia sahariensis*; Cassinello, 1997), and sorraia horses (*Equus caballus*; Heitor & Vicente, 2008). We further found that calves approached their mothers, escorts, and other females more frequently than they left them. Contrary to this, calves left their mothers more often than they approached them during the first 6 months of life in the African savannah elephant (Douglas-Hamilton 1972). However, the distance at which approaches and leavings by calves were defined in that study was 5 m, whereas we used a distance of one calf body length, which is about 1 m. Results from the African savannah elephants hint at the possibility that calves engage in different behaviours when they are within 1 m than when within 5 m of the mother, as calves were more likely to have an age mate as a neighbour in the latter scenario than in the former (Lee, 1987). This suggests that calves might engage in play when they are at a 5-m distance, resulting in them leaving their mothers more than they approach them. It is further possible that the group’s social environment – i.e., the number of individuals of different age-sex classes and the risk posed by them to the calves – may dictate conspecific female-calf approach-leave behaviour (e.g., Berman et al., 1997).

#### Behavioural interactions

Similar to proximity maintenance, we found that calves initiated and terminated most of the behavioural interactions, showing their precocial nature in forging social relationships. Similar results were reported by Gadgil and Nair (1984) in semi-captive Asian elephants (birth to 1.25 years) and by Lee (1986) in the Amboseli elephants (birth to 5 years). We found that the number of interactions initiated by calves decreased within the first six months of life, which may be a result of their increasing independence (e.g., Guarino et al., 2017; Lee, 1987). This would be consistent with the calf exploring its physical rather than social environment, as the proportion of time spent near the mother also decreased. As expected, males and females did not differ in the number of interactions initiated, consistent with previous studies (Förster & Cords, 2005; Lee, 1987), although differences between the sexes may develop with age, as found in play behaviours in the African savannah elephant (Lee & Moss, 2014).

### Calf interactions were differentially distributed among female conspecifics in the group

#### Proximity

Calves had non-random neighbours and were often positioned between their mothers and escorts, as previously observed in semi-captive Asian elephants (Gadgil & Nair, 1984). As young individuals face predation risk from tigers (Williams, 1950), being positioned near conspecific females in addition to the mother might further reduce the risk of predation (see Lee, 1987). Overall, calves showed similar proximity with mothers and escorts, but we do not have similar comparisons from other elephant populations. Newborn calves spent a greater proportion of time near their mothers than near escorts, which was also seen in semi-captive elephants (Gadgil & Nair, 1984). As the calves grew (to 3-<6 months of age), they spent more time away from their mothers; this period coincided with an increase in trunk motor skills and adult-like feeding behavioural expression (Revathe et al., 2020). Increase in skill competence has been found to be associated with an increase in mother-offspring distance in other species also (e.g., *Pongo abelii*: van Noordwijk & van Schaik, 2005). Seeking out other social partners, such as age-mates, for play, might also result in decreasing time spent near one’s mother (e.g., Arroyo-Rodríguez et al., 2007). An effect of calf age on mother-calf proximity has also been reported in wild African savannah elephants (Lee 1986) and wild and captive Asian elephants in Sri Lanka and UK facilities, respectively (Webber 2017).

We found that the proportion of time spent near escorts did not decrease significantly at least till 6 months of age. It is possible that escorts continue to actively keep track of and accompany calves as calves age, perhaps to support maternal foraging freedom while ensuring calf protection, as suggested in sperm whales (*Physeter macrocephalus*; Whitehead, 1996). Frequent care from alloparents during later stages of infant development has also been seen in other species (*Cebus olivaceus*: O’Brien & Robinson, 1991; *Canis familiaris*: Pal et al., 2021). However, our results from the Hinde’s index (albeit for calves of all ages up to 6 months) that calves were more likely to approach than to leave escorts suggest that calves may be following escorts rather than escorts moving along with calves. More data are required to differentiate between these scenarios.

The proportion of time spent in proximity to other females did not increase with calf age. Whether heterogeneity in calf-conspecific female (≥5 years) relationships is maintained as calves grow up (e.g., Berman, 1982a; Wey & Blumstein, 2010) or whether it reduces with age (Berman, 1982b; Roatti et al., 2023) remains to be seen. Overall, our findings are in line with that found in the African savannah elephant, in which mother-calf distance increased but the distance between calves and other conspecific females did not vary much with calf age (Douglas-Hamilton 1972, Lee, 1987). As expected, and in line with a previous study (Webber 2017), we did not find sex-based differences in calf-female conspecific proximity at the young ages we examined.

### Escorting females, but not others, are allomothers

#### Behavioural interactions

Despite mothers and escorts spending similar proportions of time near calves, escorts initiated more interactions (of all kinds) towards calves than did mothers. They also initiated the highest proportion of positive interactions and initiated more positive than negative interactions. Mothers also initiated more positive than negative interactions, but the difference was small. As lactation is energetically expensive and requires increased time spent foraging (Wisniewska et al., 2015) and as primarily mothers provide milk in elephants (including in the study population), other caretaking behaviours might be shared with escorts, leading to more positive interactions from escorts than mothers. A previous study in the Kabini elephant population found that females in certain clans had their first-or second-order relatives as their top associates (Nandini, 2016). If such females are escorts, they might accrue indirect fitness benefits through positive interactions with calves.

Unlike African savannah elephant calves (≤12 months old) that had been found to interact with their mothers more frequently than with any other age-sex class (Lee 1986, 1987), we found that calves (<6 months) interacted with their mothers and escorts at similarly high rates. Mothers and escorts showed similar extents of positive responses (that often required them to stop feeding) to calf-initiated interactions, terminated comparable proportions of calf-initiated interactions, and rarely directed aggression towards the calf. Mothers and escorts, but not other females, also guarded calves during long periods of calf resting. As calves <6 months old are likely in a crucial stage of learning and development, mothers and escorts may be tolerant of calf interactions through which calves explore and understand their physical and social environment and learn survival skills. Whether this relationship changes when calves attain foraging and/or social independence, as that found in great apes (Mikeliban et al., 2021) – who have learning-intensive, prolonged developmental period similar to elephants – remains to be seen.

Other females initiated fewer interactions and more negative than positive or neutral interactions than escorts did towards calves. Some of the differences in the responses of conspecifics to calves could have arisen from the fact that calves initiated similar numbers of interactions with their mothers and escorts but far fewer interactions (almost 14 times less frequent than with escorts) with other females. However, this would not explain the higher frequency of negative interactions by other females towards calves. The fact that not all the females other than the calf’s mother interacted positively – or interacted at all – with calves suggests that either conspecific females and/or the calves may actively shape their relationship, perhaps as a result of the differences in relatedness/familiarity between mothers and other conspecific females (e.g., Berman, 1982b; Dunayer & Berman, 2017). Differences in female reproductive status might also contribute to the differential nature of escort and other female interactions with calves, as females who were escorts did not have a calf of their own during the study period. However, not all the other females had a calf either during the study period.

### Calves’ role in the emergence of differentiated social relationships with female conspecifics

Calves interacted similarly with mothers and escorts in feeding, resting, and social contexts. Feeding interactions involved passive food sharing behaviours such as calves taking grass that had been scraped off the ground from a conspecific or feeding in the same patch as the conspecific. These behaviours required social tolerance and close contact between calves and conspecifics, which are prerequisites for social transmission of knowledge and skills (van Schaik, 2003), and were almost always initiated towards mothers and escorts, and almost never towards other females. Social interactions through which calves probably sought protection (from heat or predation) were also primarily with mothers and escorts, and such interactions interfered with the conspecific’s foraging, as they partly restricted their leg movement (feeding requires kicking at and scraping short grass with the foot around the Kabini backwaters). Close proximity and interactions with escorts may contribute to not just protection of young ones but also social learning opportunities in this long-lived species with a prolonged developmental period.

We found that suckling interactions accounted for about half of the calf-initiated interactions with their mothers. Calves also sucked from escorts and rarely from other females, but these were non-lactating females; thus, calves received milk only from their mothers. Calves were quickly rejected by other females the few times calves tried to suck from them, as was seen in the African savannah elephant (Lee, 1987). This is expected as lactation is energetically costly. Calves initiated more non-suckling interactions towards their escorts than even their mothers, despite similar calf-mother and calf-escort proximity. Furthermore, almost all the play interactions with conspecific females (≤5 years) were also initiated towards escorts.

We do not yet know if escorts are more closely related or more familiar to calves than other females, and whether this drives the difference in interactions. In many species, interactions of developing individuals with familiar/related individuals are more frequent than that with unfamiliar/unrelated individuals (Bădescu et al., 2015; Berman, 1982a; Konrad et al., 2019; Lee, 1987). Allomaternal care may, therefore, reinforce existing relationship differentiation. It is also possible that escorts were younger than other females, and were, therefore, preferred by calves (see Lee, 1987). Furthermore, although other females do not show overt care, they may still participate in cooperative defense of calves against predators (see McComb et al., 2011). Since our study included 20 calves from 9 elephant clans, our results are generalizable. However, we did not have enough sample size to test the effect of calf age and sex within the same model or the potential confounding effects of clan, seasonality, and year. These effects should be investigated with more data points.

In conclusion, we found that elephant groups with calves comprised females who provided active care, them being mothers and escorts, and those who did not provide overt, active care, them being other females. Females who were classified as escorts in a focal, based on coordinated movement, always provided other forms of allomaternal care during that focal. Overall, there were remarkable similarities between mothers and escorts in their behaviours towards calves and in the behaviours of calves towards them, with the primary difference being that escorts did not provide milk. These results together show that there are differentiated social relationships – i.e., variation in the quantity and quality of interactions between calves and conspecific females – from a young age. The presence of calves and their interactions with escorts in the group might increase close social interactions between mothers and escorts, which may establish new relationships or strengthen existing relationships (see also Gadgil & Nair, 1984; Lee, 1987). As calf-escort interactions involved behaviours that may potentially be helpful in feeding skill acquisition and protection, the development and survival of calves may be affected by the presence of, and/or interactions, with allomothers (Hrdy 1976, Lee 1987, Hodge 2005). The emergence of differentiated social relationships in animal societies has often been studied from the perspective of rank, sex, and age differences among the social partners of the maturing individuals, apart from differential maternal relations (including the effect of relatedness) with conspecifics. Our study highlights an additional factor - allomaternal care - which may potentially lead to the formation of new or reinforcement of existing relationship structure in animal societies. Our results, therefore, bear significance for several other mammalian study systems that show both alloparental care and differentiated adult social relationships.

## Data availability statement

The raw data will be uploaded on Dryad or the publicly available JNCASR repository after acceptance and are available immediately to the reviewers as uploaded files.

## Ethical note

Field data collection was observational in nature. Therefore, no animal handling or manipulation was involved. Researchers maintained a minimum distance of 40 m to the elephants to avoid any disturbance. Fieldwork was carried out under permit number PCCF/WL/E2/CR-121/2014-15 dated 09/05/2014, issued by the office of the Principal Chief Conservator of Forests (Wildlife) and Chief Wildlife Warden, Karnataka.

## Author contributions

Conceptualization: TR, TNCV; Data Curation: TR; Formal Analysis: TR; Funding Acquisition: TNCV; Investigation: TR; Methodology: TR, TNCV; Supervision: TNCV; Validation: TR, TNCV; Writing – Original Draft Preparation: TR; Writing – Review & Editing: TR, TNCV.

## Acknowledgements

This work was funded primarily by the Council of Scientific and Industrial Research, Government of India, under Grant No. 37(1613)/13/EMR-II, and the Jawaharlal Nehru Centre for Advanced Scientific Research (JNCASR), and in part by the Science and Engineering Research Board (SERB), Government of India, under Grant No. CRG/2022/005975. TR was supported as a student by JNCASR. This work is part of TR’s Ph.D. thesis. We thank the offices of the PCCF(WL) and APCCF(WL), Karnataka Forest Department, and of the Conservators of Forests of Nagarahole and Bandipur National Parks and Tiger Reserves for field permits. We also thank other officials and staff of Nagarahole and Bandipur National Parks for their support. We thank Ranga, Krishna, Shankar, Pramod, and others for field assistance. We thank Hansraj Gautam for recording some focal calf videos and members of the Animal Behaviour Lab, especially Ankana Sanyal, Anvitha S, Athira TK, Divya Choudhary, and Jabili Chowdari, for help with checking field data. We thank Amitabh Joshi for suggestions on statistical analyses.

## Supplementary Material

Supplementary Material 1. Details of focal calves.

**Supplementary Material 1, Table 1.** Details of focal calves including their clan identities, age-class (at the time of sampling), and sex that were sampled to get durations of calf positions, calf proximity, and calf behavioural interactions data.

| S.No. | Calf identity | Clan identity | Calf sex | Calf age-class |
| --- | --- | --- | --- | --- |
| 1 | Genette_2017_F | Patricia | F | <3 months |
| 2 | Georgina_2017_F | Patricia | F | <3 months |
| 3 | Ilaena_2016_F | Menaka | F | <3 months |
| 4 | Alena_2017_M | Alexandra | M | <3 months |
| 5 | Amarilla_2016_M | Osanna | M | <3 months |
| 6 | Gemini_2017_M | Patricia | M | <3 months |
| 7 | Iliora_2015_M | Menaka | M | <3 months |
| 8 | Ipomoea_2018_M | Victoria | M | <3 months |
| 9 | Kai_2016_M | Nakshatra | M | <3 months |
| 10 | Kasturi_2018_M | Kasturi | M | <3 months |
| 11 | Floppy_ears_2018_F | Victoria | F | 3-<6 months |
| 12 | Marlene_2015_F | Tilottama | F | 3-<6 months |
| 13 | Namrata_2017_F | Nakshatra | F | 3-<6 months |
| 14 | Suhrita_2016_F | Victoria | F | 3-<6 months |
| 15 | Zarin_2015_F | Victoria | F | 3-<6 months |
| 16 | Camila_2017_M | Patricia | M | 3-<6 months |
| 17 | Kausalya_2015_M | Kasturi | M | 3-<6 months |
| 18 | Orlanda_2015_M | Lisa | M | 3-<6 months |
| 19 | Valerie_2016_M | Victoria | M | 3-<6 months |
| 20 | Vanessa_2015_M | Victoria | M | 3-<6 months |

**Supplementary Material 1, Table 2.** Details of focal calves, including their clan identities and age-, sex-class that were sampled to get durations of calf positions to calculate the time to independence of calf positions.

| Calf identity | Clan identity | Calf sex | Calf age (in months) | Calf age-class | No. of different calf positions |
| --- | --- | --- | --- | --- | --- |
| Kausalya_2015_M | Kasturi | M | 4.33 | <6 months | 25 |
| Kausalya_2015_M | Kasturi | M | 4.43 | <6 months | 12 |
| Kausalya_2015_M | Kasturi | M | 4.97 | <6 months | 18 |
| Hannah_2016_F | Tilottama | F | 3.80 | <6 months | 16 |
| Hannah_2016_F | Tilottama | F | 3.97 | <6 months | 20 |
| Hannah_2016_F | Tilottama | F | 4.63 | <6 months | 22 |
| Vanessa_2015_M | Victoria | M | 5.23 | <6 months | 6 |
| Vanessa_2015_M | Victoria | M | 5.73 | <6 months | 27 |
| Vanessa_2015_M | Victoria | M | 5.97 | <6 months | 45 |
| Genette_2017_F | Patricia | F | 0.10 | <6 months | 22 |
| Genette_2017_F | Patricia | F | 1.00 | <6 months | 8 |
| Genette_2017_F | Patricia | F | 1.17 | <6 months | 22 |
| Leena_2017_F | Lisa | F | 3.37 | <6 months | 15 |
| Leena_2017_F | Lisa | F | 3.60 | <6 months | 9 |
| Leena_2017_F | Lisa | F | 4.00 | <6 months | 32 |

Supplementary Material 2. Calf position codes and duration to independence of positions.

**Supplementary Material 2, Table 1.** Focal calf’s position codes and their descriptions. Mother refers to the focal calf’s mother; escort/s refers to the focal calf’s escort/s; other female/s refers to females (≥ 5 years) other the focal calf’s mother or escorts in a group; conspecific male refers to males (≥ 5 years) in a group; juvenile refers to a female or a male juvenile in the group; and another calf refers to a female or a male calf other than the focal calf in the group. Near refers to the focal calf standing/sitting/lying down within one calf body length of any of the conspecifics. The order in which conspecific categories (Calf/Juvenile/Mother/Escort/Other female/Male) appear in the position code do not signify anything.

| S.No. | Focal calf position code | Code description |
| --- | --- | --- |
| 1 | ALO | Alone |
| 2 | SNC | Near another calf |
| 3 | SCE | Near another calf and an escort |
| 4 | CCJ | Near another calf and a juvenile |
| 5 | FCJ | Near another calf, a juvenile, and an other female |
| 6 | SCM | Near another calf and the mother |
| 7 | CCC | Near another calf and two other females |
| 8 | SNE | Near an escort and not near any other individual |
| 9 | ESC | Near an escort and another calf |
| 10 | SEE | Near two escorts |
| 11 | SEC | Near an escort and an other female |
| 12 | EFC | Near an escort, an other female, and another calf |
| 13 | EFJ | Near an escort, an other female, and a juvenile |
| 14 | ECC | Near an escort and two other females |
| 15 | SNJ | Near a juvenile |
| 16 | SJE | Near a juvenile and an escort |
| 17 | SJJ | Near two juveniles |
| 18 | SJM | Near a juvenile and the mother |
| 19 | SML | Near a conspecific male |
| 20 | SME | Near a conspecific male and an escort |
| 21 | SMM | Near a conspecific male and the mother |
| 22 | SMF | Near a conspecific male and an other female |
| 23 | MFE | Near a conspecific male, an other female, and an escort |
| 24 | SNM | Near the mother and not near any other individual |
| 25 | SEM | Near an escort and the mother |
| 26 | MEC | Near the mother, an escort, and another calf |
| 27 | MEE | Near the mother and two escorts |
| 28 | MEJ | Near the mother, an escort, and a juvenile |
| 29 | MEM | Near the mother, an escort, and a conspecific male |
| 30 | MEF | Near the mother, an escort, and an other female |
| 31 | SMC | Near the mother and an other female |
| 32 | MFC | Near the mother, an other female, and another calf |
| 33 | MFJ | Near the mother, an other female, and a juvenile |
| 34 | MCC | Near the mother and two other females |
| 35 | MCO | Near the mother, two other females, and a calf |
| 36 | SCO | Near an other female and not near any other individual |
| 37 | SFC | Near an other female and another calf |
| 38 | FEE | Near an other female and two escorts |
| 39 | SFJ | Near an other female and a juvenile |
| 40 | SCC | Near two other females |

While a focal calf could be near other calves or juveniles or subadult/adult males (as in the table above), as we were not interested in looking at these contacts, we did not include them in the analysis. Only changes in calf positions when alone or involving conspecific females were considered for the analysis.

### Durations of calf positions

We constructed cumulative frequency distributions of durations of calf positions, calculated from second-to-second scoring of calf position changes for calves <6 months of age. Position changes were included irrespective of whether the calf or the conspecific moved to bring about the change. We found that 95% of the calf positions (near conspecific females ≥5 years or alone) changed within 200-210 seconds for calves <6 months (see figure below). Therefore, we considered two subsequent calf positions to be independent if they were separated by 4 minutes.

**Supplementary Material 2, Figure 1.**
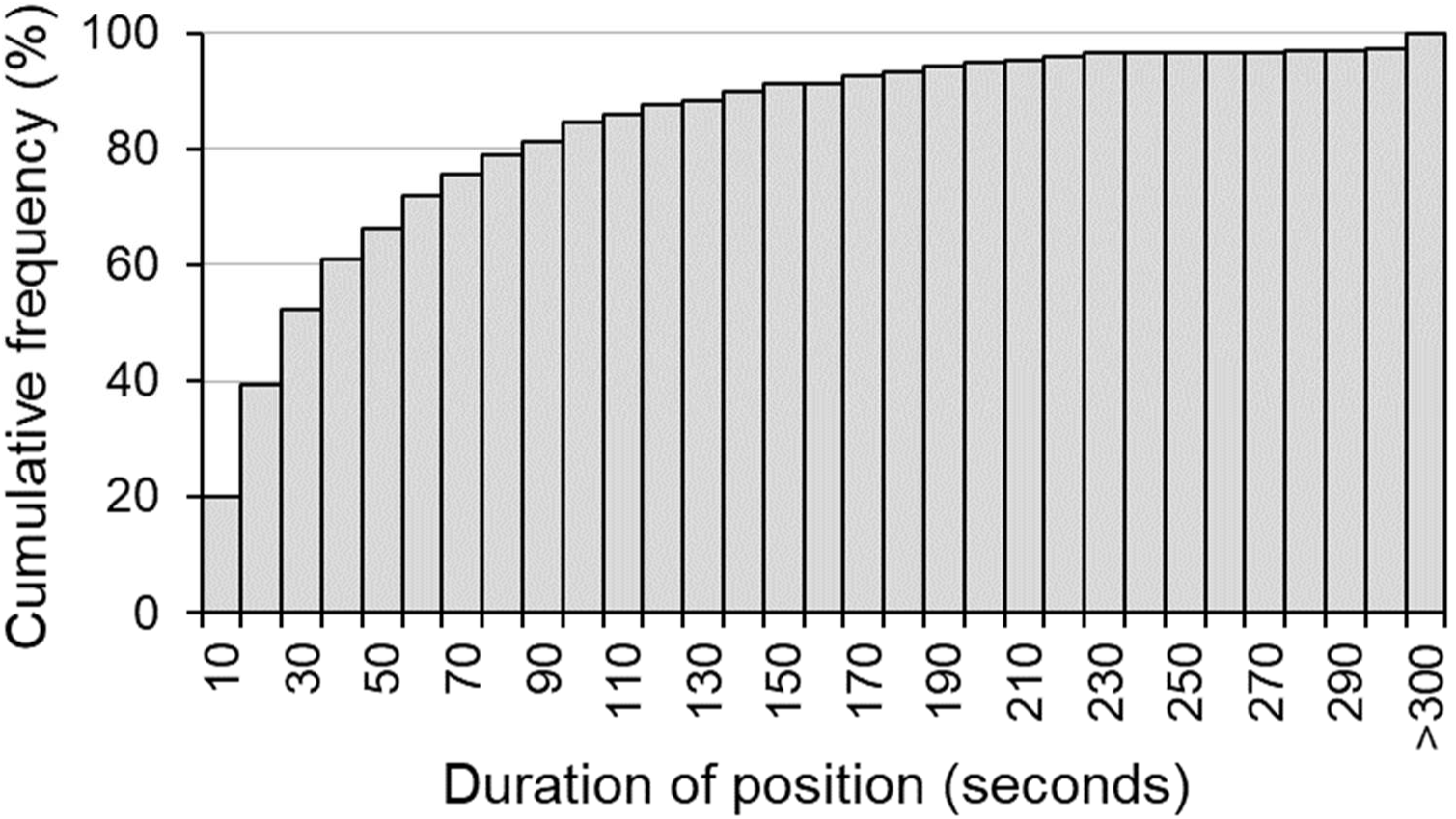
Cumulative frequency distributions of the time durations of calf positions for calves that were <6 months of age.

Supplementary Material 3. Details of Hinde’s and Brown’s proximity indices.

We examined calf-conspecific proximity initiations using Hinde’s proximity index and Brown’s proximity index.

### Hinde’s proximity index

Hinde’s proximity index (Hinde and Spencer-Booth 1967) was originally developed as a measure to understand the dynamics of mother-infant relationships in primates. Whenever the calf’s position changed, we recorded the time and whether the change was due to the calf’s or conspecific’s movement. An approach (*Ap*) was recorded when the distance between a calf-conspecific pair decreased from >1 to ≤1 calf-body length (either due to the calf’s action – *Ap_c_* or conspecific’s action – *Ap_cf_*), while a leaving (*L*; *L_c_* when a calf left, and *L_cf_* when conspecific left) was recorded when the calf-conspecific distance increased from ≤1 to >1 calf-body length. Hinde’s proximity index for each calf-conspecific pair was calculated as follows to see whether calves showed more approaches or leavings towards each conspecific category:

Hinde’s index = the percentage of approaches (% *Ap_c_*) made by a young one (calf here) towards a conspecific – the percentage of leavings made by that young one away from that conspecific (% *L_c_*).

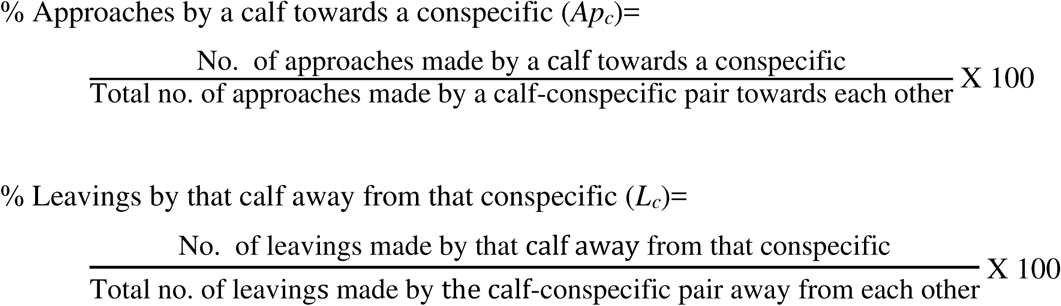

The index indicates whether the calf more frequently approached or left a particular conspecific. Hinde’s proximity index varies between -100 to +100 (Figure). A value of 0 indicates that the young one makes and breaks proximity contacts with a conspecific equally; a negative value indicates that the calf more often leaves than approaches the conspecific, and a positive value that the calf more often approaches than leaves the conspecific. While the total number of approaches would be similar to the total number of leavings (*Ap* = *L* (±1) in a focal) for a calf-conspecific pair (Hinde and Atkinson 1970), the number of approaches by the calf (*Ap_c_*) could differ from the number of leavings by the calf (*L_c_*) for the pair.

### Brown’s proximity index

Hinde’s index does not measure the relative contributions of the calf and conspecific to changes in proximity contacts between the pair. Therefore, we also the calculated Brown’s proximity index (Brown, 2001) to measure the relative contributions of the calf and conspecific to changes in proximity contacts between the pair. This calculates the total percentage of changes in proximity contacts that were due to the movement of the calf.

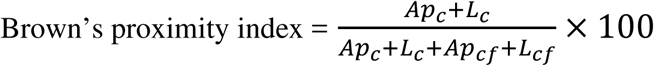

where,

*Ap_c_* and *L_c_* are as above,

*Ap_cf_* is the number of approaches made by the conspecific towards the calf

*L_cf_* is the number of leavings made by the conspecific away from the calf.

**Supplementary Material 3, Figure 1.**
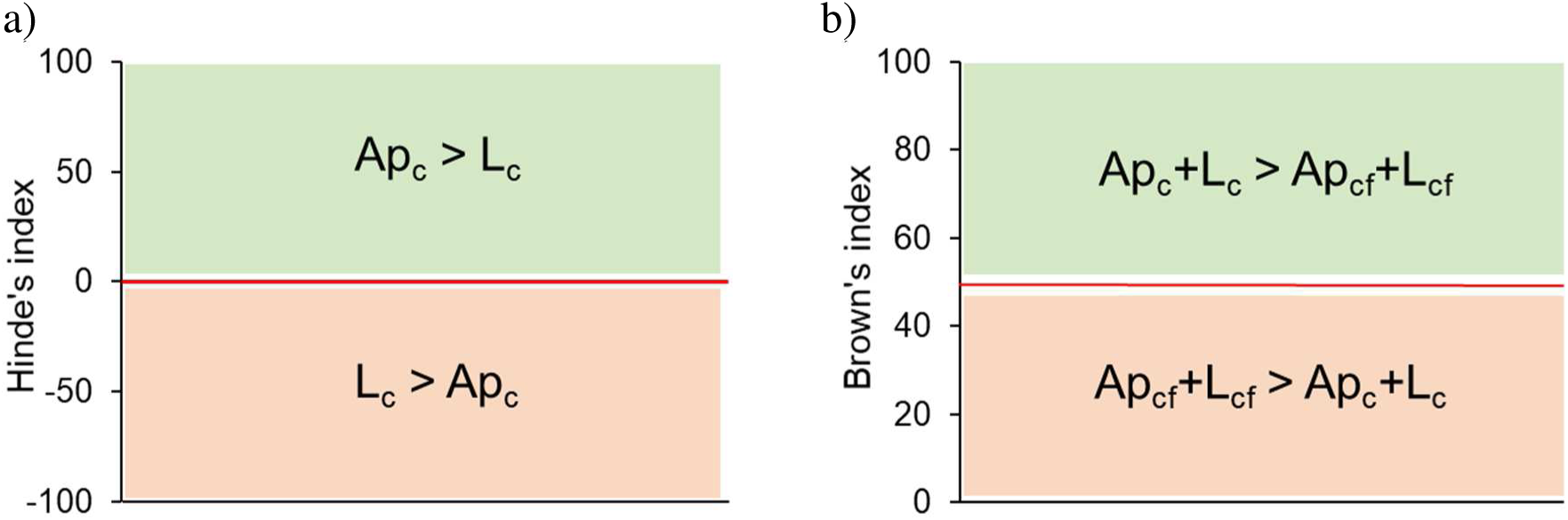
A schematic of a) Hinde’s and b) Brown’s proximity indices.

Brown’s index ranges from 0 to 100, with 0 indicating that the calf was not responsible for any of the changes in proximity (numerator will be zero), a value of 50 indicating that young one and conspecific were equally responsible for changes in proximity contacts, any value between 0 and 50 indicating that the conspecific was more responsible for changes in proximity contacts than the young one, a value of 100 indicating that the calf was responsible for all the changes in proximity, and any value between 50 and 100 indicating that the calf was more responsible for changes in proximity contacts than the conspecific. While Brown’s proximity index measures the relative contribution of calves and conspecifics to the changes in proximity contacts, it does not differentiate between calves approaching or leaving a conspecific to a greater extent. Hinde’s and Brown’s proximity indices complement each other by indicating whether the calf was responsible for making or breaking contact with a particular conspecific and whether the calf or conspecific contributed more to the changes in proximity between them. The values of both the indices may change with the identity of the conspecific (Rowell *et al*. 1964) and the young one’s age (Rowell *et al*. 1964, Hinde and Spencer-Booth 1967, Hinde and Atkinson 1970, Douglas-Hamilton 1972), and the indices are not correlated (Brown 2001). The relative contributions of the young one versus the conspecific in proximity contact changes would also differ based on the distance at which approaches and leavings were measured (Brown 2001).

If a focal calf had more than one escort within a focal sample, the Hinde’s index and Brown’s index were averaged across escorts. Similarly, averages were used when a focal calf had more than one ‘other female’ with which it was involved in a proximity initiation/termination. As mentioned in the main text, in certain focals, despite the presence of three conspecific categories of females in a focal for a focal calf, there was no proximity initiation/termination between the calf and one or more categories of females. Therefore, average Hinde’s and Brown’s proximity indices could be calculated only for 7 calves (*N*: 3 females, 4 males; 3 newborn calves (<3 months), 4 infant calves (3-<6 months)) that each had proximity initiation/termination with all three conspecific categories of females in at least two of their focals. There was no significant difference in the Hinde’s (Friedman ANOVA: χ^2^= 2.000, *N*=7, *df*=2, *P*=0.368) or Brown’s (Friedman ANOVA: χ^2^= 4.571, *N*=7, *df*=2, *P*=0.102) proximity indices among mothers, escorts, and other females. ANOVAs on the logit Hinde’s index with mothers and escorts based on data from a larger number of calves (17 calves, 44 focals) also showed no significant effect of conspecific category (*F*_1,16_=3.849, *P*=0.067). There was no effect of calf ID (*F*_16,54_=0.992, *P*=0.479) or its interaction with conspecific category in the ANOVA (*F*_16,54_=0.724, *P*=0.757), but inferences cannot be drawn based on these as each calf was sampled only twice. ANOVA on the logit Brown’s index with mothers and escorts (17 calves, 44 focals) also showed no significant effect of conspecific category (*F*_1,16_=0.888, *P*=0.360), calf ID (*F*_16,54_=1.484, *P*=0.140), or their interaction (*F*_16,54_=0.365, *P*=0.985).

Supplementary Material 4. Details of behavioural interactions recorded.

As mentioned in the main text, we used focal animal sampling (Altmann, 1974) to record interactions between calves and conspecifics within groups. We recorded observations using a video camera (SONY HDR-XR 100E or SONY HDR-PJ 540E). Focal videos were taken in such a way that a focal calf was always in frame to capture all the calf-conspecific (i.e., subadult or adult female) interactions that occurred. We also ensured that the focal calf’s mother was also in the frame, except in cases when the calf ventured far away from its mother, in which case, the mother’s behaviours were written down. Calf-initiated interactions, conspecific-initiated interactions, and responses by conspecifics are detailed in the tables below.

**Supplementary Material 4, Table 1.** Calf-initiated interactions towards mother, escorts, and other females, the frequencies with which they appear in the data, behavioural classes, and descriptions of the interactions.

| Code | Interaction | Frequency | Behavioural class | Description |
| --- | --- | --- | --- | --- |
| INV | Investigate food | 38 | Feeding | Investigate another animal's food (in its mouth)/smell the grass that an individual is scraping off, but NOT taking someone else's food (which is TGC). |
| NDG | Nudge | 1 | Feeding | Nudge another animal especially when competing for food and feeding very close by and almost always supplanting the animal and feeding there (not dominance related in case of calves). |
| NPL | Pull at nipple | 5 | Feeding | Pull or hold nipple. |
| NPR | Locate nipple correctly | 192 | Feeding | Try to locate nipple in the correct direction. This includes sniffing in the direction of the nipples also. |
| NPW | Search for nipple in the wrong direction | 9 | Feeding | Try to locate nipple between the hind legs (also includes trying to suck from there). |
| SUA | Allosuck | 45 | Feeding | Suck from the escort/allomother. |
| SUF | Suck from female | 2 | Feeding | Suck from a female conspecific other than the mother or escort. |
| SUM | Suck from mother | 137 | Feeding | Suck from mother. |
| TGC | Take grass from a conspecific | 89 | Feeding | Take grass from a conspecific (and feed on it; PLO if they just throw it away PLO). |
| TGS | Take grass from a spot | 21 | Feeding | Feed from the exact spot where another is feeding from (mostly with mothers/escorts). |
| LEN | Lean | 27 | Resting | Stand or sit leaning on another animal. |
| LIE | Lie down | 54 | Resting | Lie down on land (as opposed to lying in the water, which is bathe BTH). |
| SLD | Slide | 8 | Resting | Slide off an individual's leg to lie down. |
| CHE | Bite | 1 | Social | Try to chew or bite any body part of another individual. |
| CHK2 | Check | 12 | Social | Check a conspecific but NOT in dominance. |
| DST | Express distress | 3 | Social | Express distress through vocalisations. |
| EXT | Extend trunk | 1 | Social | Extend trunk towards a conspecific. |
| KIC | Kick conspecific | 1 | Social | Kick a conspecific, even if it misses. |
| PCL | Play climb | 2 | Social | Climb/roll on other calves (play). |
| PSH | Push | 16 | Social | Push with the head. |
| PTS | Pass | 36 | Social | Pass from one side of an older animal to the other through the space between that animal's trunk and forelegs or through the space under its belly. |
| RUB | Rub | 151 | Social | Rub against a conspecific (no dominance). |
| SBE | Stand with<br>body contact | 46 | Social | Stand between the trunk and leg of an escort. |
| SBM | Stand with<br>body contact | 11 | Social | Stand between the trunk and leg of the mother. |
| SNI | Sniff air | 62 | Social | Sniff the air (could be in the direction of a conspecific or heterospecific). |
| STE | Stand with<br>escort body<br>contact | 46 | Social | Stand under the escort |
| STM | Stand with<br>mother body<br>contact | 39 | Social | Stand under the mother. |
| TOU | Touch | 129 | Social | Touch with trunk in areas apart from the mouth (in mouth would be TRM) and genitals (genitals would be CHK or CHQ), and not in dominance (dominance would be TCH). |

**Supplementary Material 4, Table 2.**
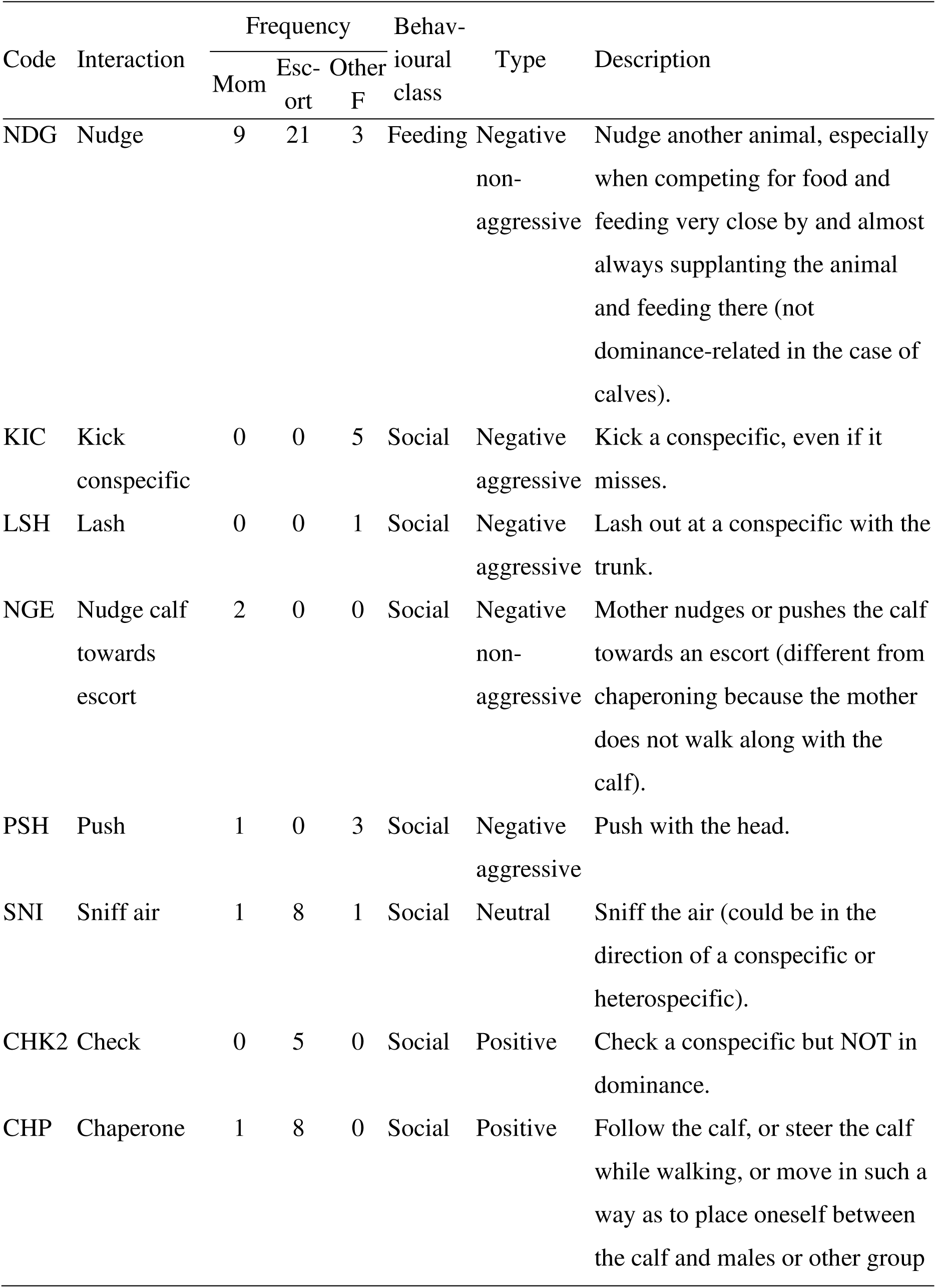
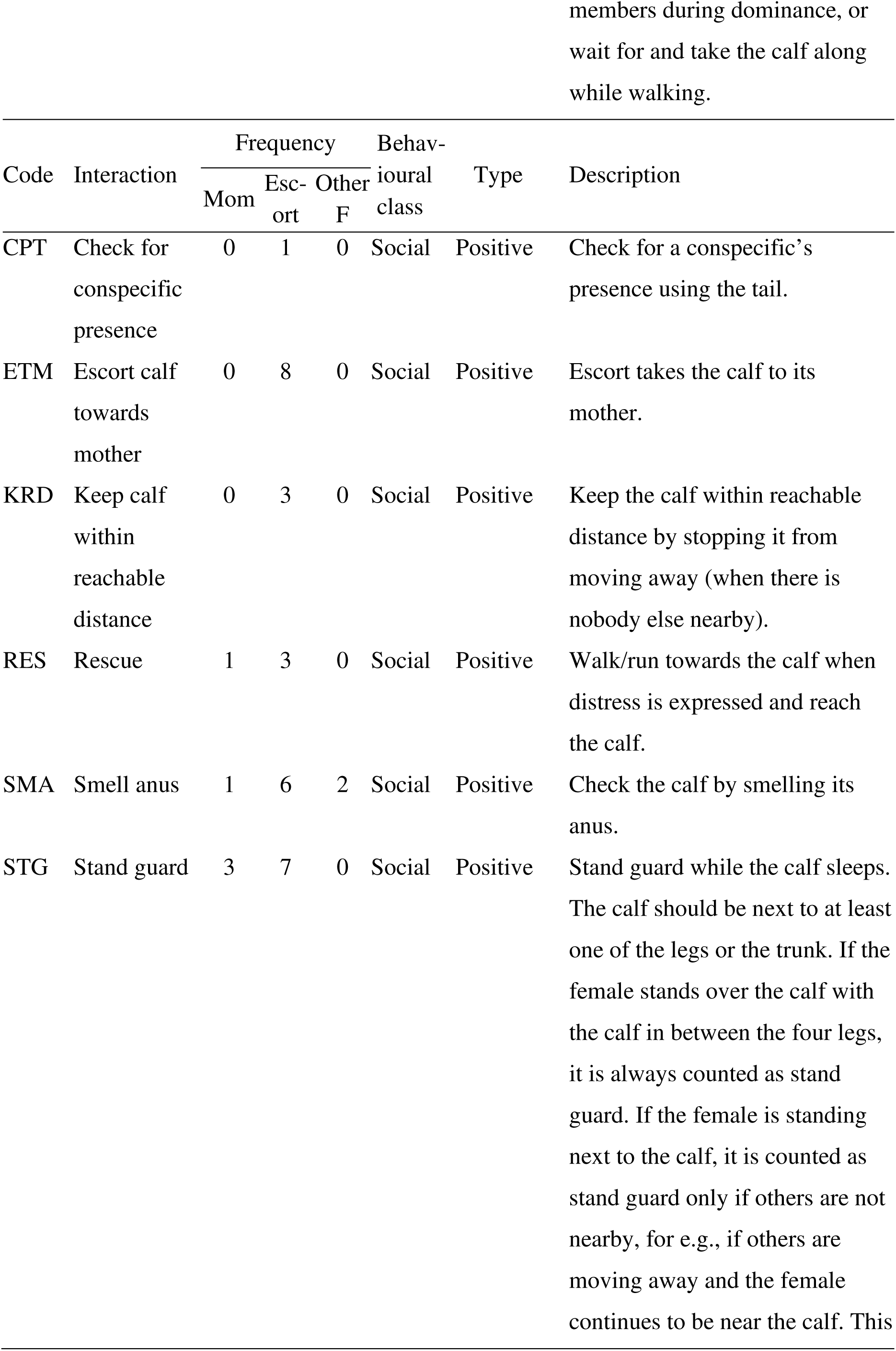
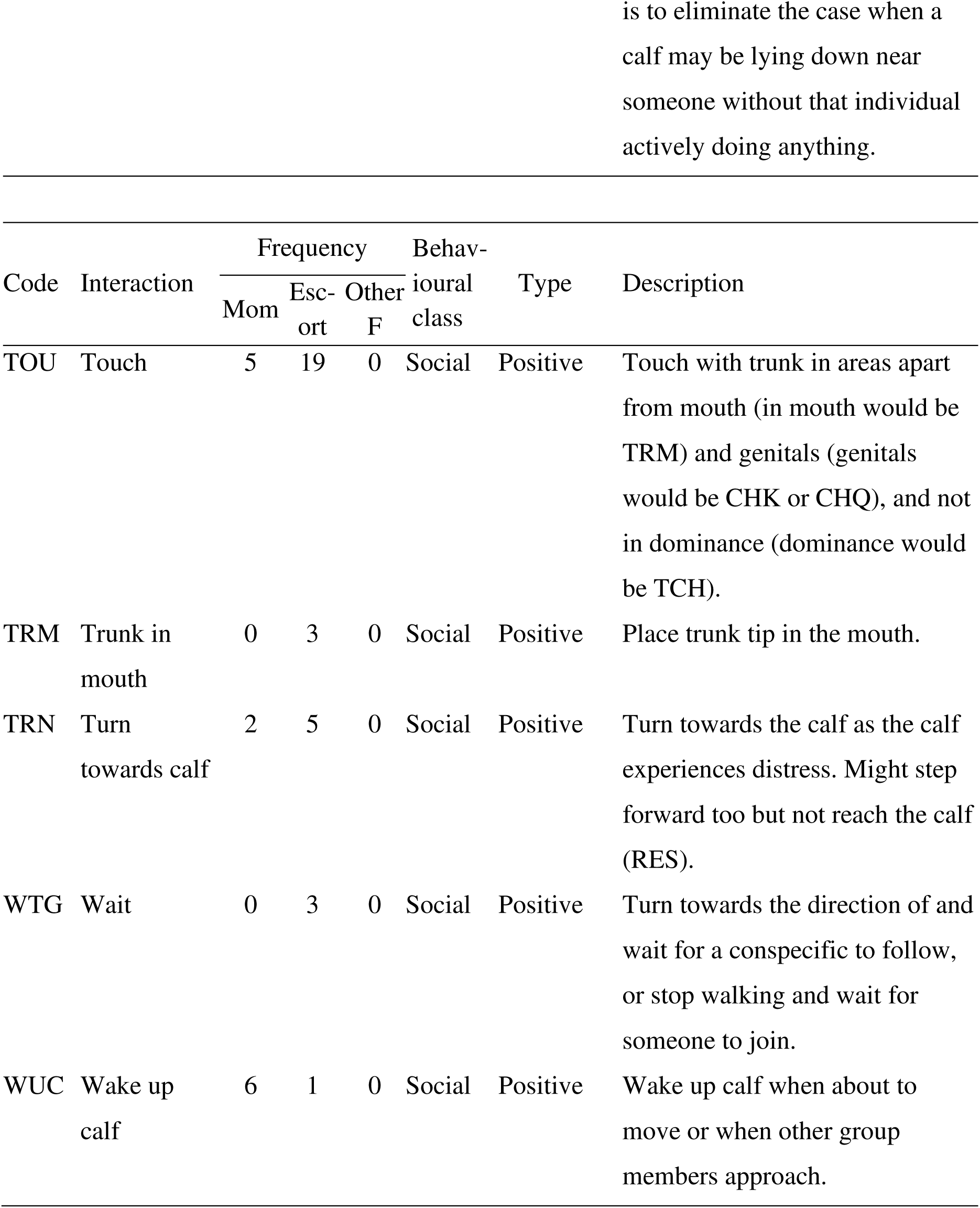
Conspecific-(mother, escorts, and other females) initiated interactions towards calves, the frequencies with which they appear in the data, behavioural classes, type, and descriptions of the interactions.

It must be noted that as in the case of proximity, it was possible, although uncommon, for more than one conspecific to simultaneously interact with the same calf.

**Supplementary Material 4, Table 3.** Responses received by the calves to the interactions initiated towards mothers, escorts, and other females, the frequencies with which these responses appear in the data, and the type and the descriptions of the responses.

| Code | Response | Frequency | Type | Description |
| --- | --- | --- | --- | --- |
| AVO | Avoid | 38 | Negative | Turn away/walk away. |
| KIC | Kick | 4 | Negative | Kick a conspecific even if it misses. |
| LSH | Lash | 4 | Negative | Lash out at a conspecific with trunk. |
| MVL | Move leg | 99 | Negative | Move leg to stop calf from rubbing/leaning/sucking. |
| NDG | Nudge | 13 | Negative | Nudge another animal especially when competing for food and feeding very close by and almost always supplanting the animal and feeding there (not dominance related in the case of calves). |
| PSH | Push | 5 | Negative | Push with the head. |
| PTR | Pull trunk | 1 | Negative | Pull (hold) the trunk and stop the individual from feeding. |
| SFT | Shift | 25 | Negative | Shift body/trunk but not move away from the spot; adjust one's position in such a way that it stops the calf's body contact with the recipient. |
| TLB | Beat with tail | 3 | Negative | Beat a conspecific with one's tail. |
| NOR | No response | 884 | Neutral | No response. |
| SNI | Sniff air | 1 | Neutral | Sniff the air (could be in the direction of a conspecific or heterospecific). |
| ATN | Pay attention | 3 | Positive | Stand next to the calf and watch it (do something); the individual has to stop whatever it is doing and pay attention to the calf. |
| CCA | Cooperate with calf | 14 | Positive | Cooperate with calf action by stopping what one is doing. |
| CSU | Cooperate with suckling | 62 | Positive | Move foreleg forward or stop feeding for the calf to suck, or stand without moving once the calf starts sucking. |
| RES | Rescue | 1 | Positive | Walk/run towards the calf when distress is expressed and reach the calf. |
| SMA | Smell<br>anus | 1 | Positive | Check the calf by smelling the anus. |
| STG | Stand<br>guard | 16 | Positive | Stand guard while the calf sleeps. The calf should be next to at least one of the legs or the trunk. If the female stands over the calf with the calf in between the four legs, it is always counted as stand guard. If the female is standing next to the calf, it is counted as stand guard only if others are not nearby, for e.g., if others are moving away and the female continues to be near the calf. This is to eliminate the case when a calf may be lying down near someone without that individual actively doing anything. |
| TOU | Touch | 2 | Positive | Touch with trunk in areas apart from mouth (in mouth would be TRM) and genitals (genitals would be CHK or CHQ), and not in dominance (dominance would be TCH). |
| TRM | Trunk in<br>mouth | 4 | Positive | Place trunk tip in the mouth. |
| WUC | Wake up<br>calf | 4 | Positive | Wake up calf when about to move or when other group members approach. |

Supplementary Material 5. Effect of calf sex on calf-conspecific proximity.

**Supplementary Material 5, Table 1.** Results of the nested ANOVA on the logit proportion of scans that calves spent near the three conspecific categories of females using 5 female and 5 male infant calves (3-<6 months old). Significant *P* values are marked in bold.

| Effect | Effect<br>(F/R) | *SS* | *df* | *MS* | *F* | *P* |
| --- | --- | --- | --- | --- | --- | --- |
| Calf sex | Fixed | 10.354 | 1 | 10.354 | 2.956 | 0.124 |
| Conspecific category | Fixed | 352.577 | 2 | 176.289 | 32.175 | **<0.001** |
| Calf sex*Conspecific category | Fixed | 2.358 | 2 | 1.179 | 0.215 | 0.809 |
| Calf ID (calf sex) | Random | 28.020 | 8 | 3.502 | 0.942 | 0.490 |
| Calf ID (calf sex)*Conspecific category | Random | 87.665 | 16 | 5.479 | 1.473 | 0.140 |
| Error | — | 223.134 | 60 | 3.719 | — | — |

Note: Inferences should not be drawn about Calf ID or interactions with this factor as each calf was sampled in only two focals.
Note: As a calf could have more than one escort and/or other female in a focal, we calculated the proportions of scans that the focal calf spent near each of the three conspecific categories of females as the sum of the number of scans near each of the females in that conspecific category / (total number of females in that conspecific category x 6; where 6 was the number of total scans in each focal). Since calves could simultaneously be near more than one category of conspecifics sometimes, the sum of the three proportions could exceed 1.

Supplementary Material 6. Durations of calf-conspecific interactions.

**Supplementary Material 6, Figure 1.**
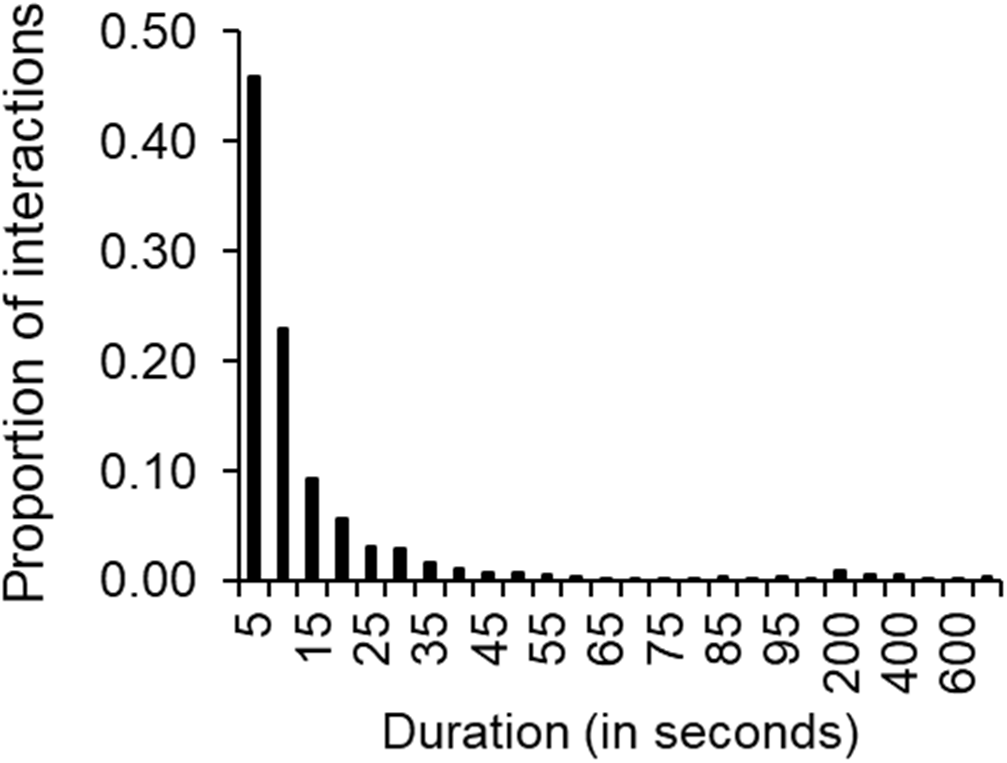
Frequency distribution of the durations of calf-conspecific interactions (initiated by either the calf or the conspecific).

Analyses carried out on the durations of interactions (variables: calf-conspecific interactions and calf-initiated interactions under different behavioural classes) gave similar results as those on the numbers of calf-conspecific interactions and numbers of calf-initiated interactions under different behavioural classes. Therefore, they are not reported here.

Supplementary Material 7. Effect of calf sex on the numbers of calf-conspecific interactions.

We ran a nested ANOVA with the log-transformed numbers of calf-conspecific interactions as the dependent variable, calf sex, initiator category (Calf, Conspecific), and conspecific category (Mother, Escort, and Other F) as fixed factors, and calf ID nested within sex as a random factor. We also checked the interaction effects. We could not test the effect of sex for newborn calves, as there were only 3 female (and 7 male) calves. We ran a nested ANOVA with only infant calves (3-<6 months), which contained 5 female and 5 male calves. The effect of calf sex was not significant. However, even these are small sample sizes and it is desirable to test the effects of calf sex on proximity and behavioural interactions with larger sample sizes in the future.

**Supplementary Material 7, Table 1.** Results of the nested ANOVA on the log no. of all calf-conspecific interactions using 5 female and 5 male infant calves (3-<6 months).

| Effect | Effect (F/R) | SS | df | MS | F | P |
| --- | --- | --- | --- | --- | --- | --- |
| Calf ID(calf sex) | Random | 7.209 | 8 | 0.901 | 2.195 | **0.040** |
| Calf sex | Fixed | 0.074 | 1 | 0.074 | 0.082 | 0.782 |
| Initiator category | Fixed | 58.964 | 1 | 58.964 | 133.247 | **<0.001** |
| Conspecific category | Fixed | 27.505 | 2 | 13.753 | 35.559 | **<0.001** |
| Calf ID(calf sex)*Initiator category | Random | 3.540 | 8 | 0.443 | 1.078 | 0.391 |
| Calf ID(calf sex)*Conspecific category | Random | 6.188 | 16 | 0.387 | 0.942 | 0.528 |
| Calf sex*Initiator category | Fixed | 0.333 | 1 | 0.333 | 0.753 | 0.411 |
| Calf sex*Conspecific category | Fixed | 0.313 | 2 | 0.156 | 0.404 | 0.674 |
| Initiator category*Conspecific category | Fixed | 20.936 | 2 | 10.468 | 37.391 | **<0.001** |
| Calf ID(calf sex)*Initiator category*Conspecific category | Random | 4.479 | 16 | 0.280 | 0.682 | 0.800 |
| Calf sex*Initiator category*Conspecific category | Fixed | 0.204 | 2 | 0.102 | 0.365 | 0.700 |
| Error | — | 24.627 | 60 | 0.410 | — | — |

Note: Inferences should not be drawn about Calf ID or interactions with this factor as each calf was sampled in only two focals.

Supplementary Material 8. Non-suckling interactions.

Of the 1184 calf-initiated interactions, there was a total of 794 non-suckling interactions and 390 suckling interactions towards mothers, escort, and other females. The nested ANOVA on the number of non-suckling interactions showed almost the same pattern of results as that on all the interactions. There was a significant main effect of calf age-class, initiator category, and conspecific category on the log number of non-suckling calf-conspecific interactions (Table 1 below). Newborn calves initiated a significantly higher number of non-suckling interactions with conspecific females than infant calves (Figure 1 below). Even after removing suckling interactions, calves continued to initiate a significantly higher number of interactions (average ± 95% CI) towards conspecific females (19.85 ± 3.770) than the latter did towards calves (3.70 ± 1.634). There was again a significant interaction effect between initiator category and conspecific category (Table 1 below). Calves initiated a significantly higher number of non-suckling interactions towards their mothers and escorts than towards other females (*P*<0.05 for both the comparisons), whereas escorts initiated a higher number of interactions towards calves than mothers and other females did (*P*<0.05 for both the comparisons). Unlike the case of all interactions, calves initiated more non-suckling interactions towards escorts than towards their mothers (*P*<0.05). There was a significant effect of calf identity and also its interaction with conspecific category, but the other random factors were not significant.

**Supplementary Material 8, Table 1.**
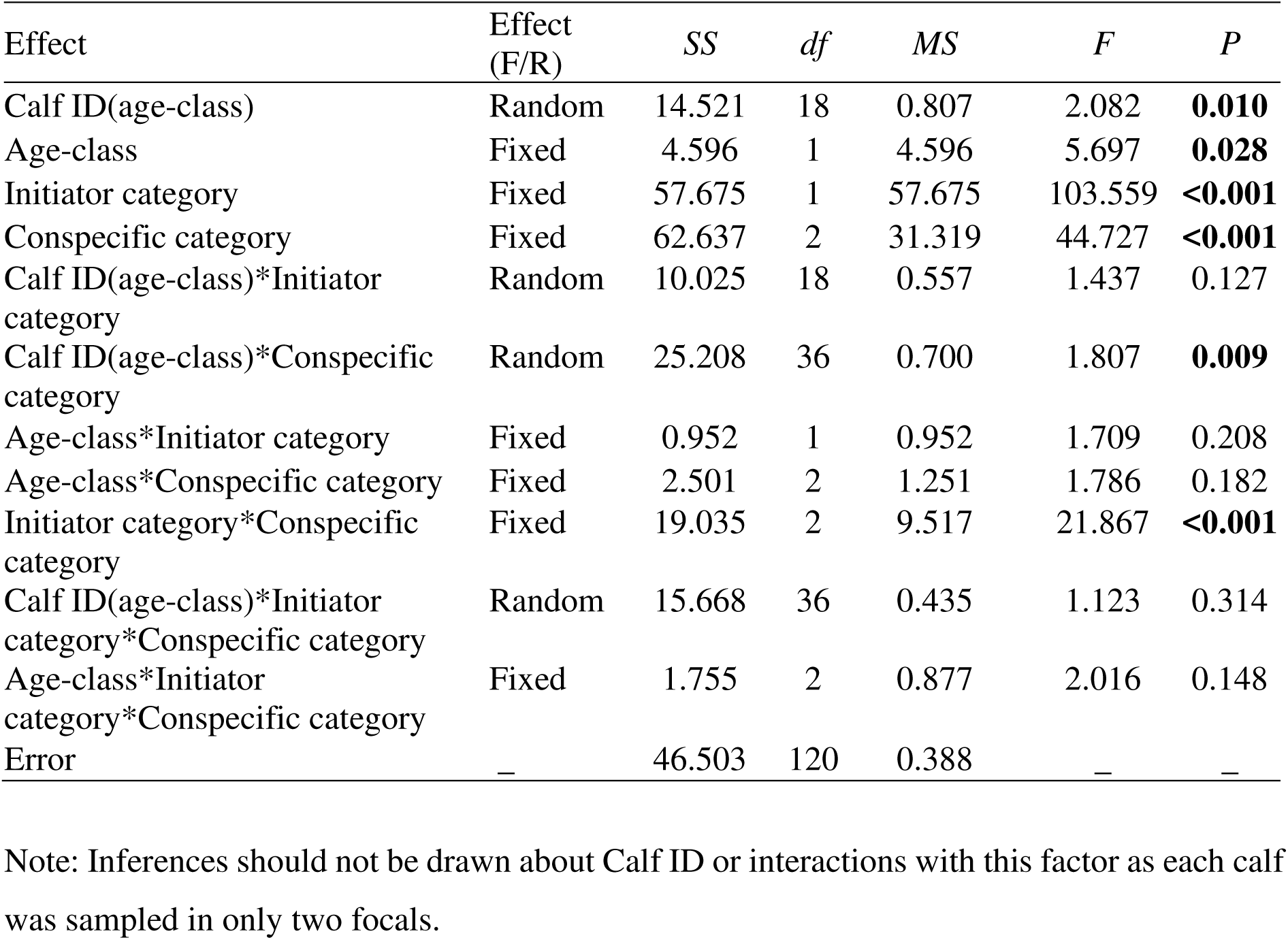
Results of the nested ANOVA on the log numbers of non-suckling calf-conspecific interactions. Significant *P* values are marked in bold.

| Effect | Effect (F/R) | *SS* | *df* | *MS* | *F* | *P* |
| --- | --- | --- | --- | --- | --- | --- |
| Calf ID(age-class) | Random | 14.521 | 18 | 0.807 | 2.082 | **0.010** |
| Age-class | Fixed | 4.596 | 1 | 4.596 | 5.697 | **0.028** |
| Initiator category | Fixed | 57.675 | 1 | 57.675 | 103.559 | **<0.001** |
| Conspecific category | Fixed | 62.637 | 2 | 31.319 | 44.727 | **<0.001** |
| Calf ID(age-class)*Initiator category | Random | 10.025 | 18 | 0.557 | 1.437 | 0.127 |
| Calf ID(age-class)*Conspecific category | Random | 25.208 | 36 | 0.700 | 1.807 | **0.009** |
| Age-class*Initiator category | Fixed | 0.952 | 1 | 0.952 | 1.709 | 0.208 |
| Age-class*Conspecific category | Fixed | 2.501 | 2 | 1.251 | 1.786 | 0.182 |
| Initiator category*Conspecific category | Fixed | 19.035 | 2 | 9.517 | 21.867 | **<0.001** |
| Calf ID(age-class)*Initiator category*Conspecific category | Random | 15.668 | 36 | 0.435 | 1.123 | 0.314 |
| Age-class*Initiator category*Conspecific category | Fixed | 1.755 | 2 | 0.877 | 2.016 | 0.148 |
| Error | — | 46.503 | 120 | 0.388 | — | — |

Note: Inferences should not be drawn about Calf ID or interactions with this factor as each calf was sampled in only two focals.

**Supplementary Material 8, Figure 1.**
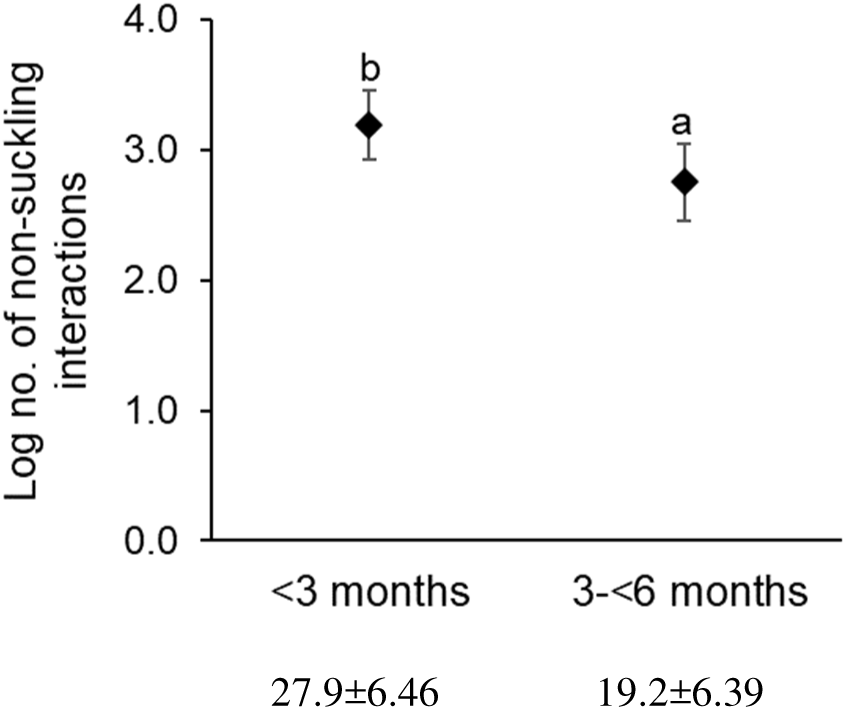
Log numbers of non-suckling interactions between newborn and infant calves and conspecific females. Error bars are 95% CI. Letters above the data points indicate the pattern of statistical significance based on Tukey’s HSD tests (a<b). The values of the number of interactions (average ± 95% CI) are shown below the graphs.

Supplementary Material 9. Details of focals that had calf-conspecific interactions.

**Supplementary Material 9, Table 1.**
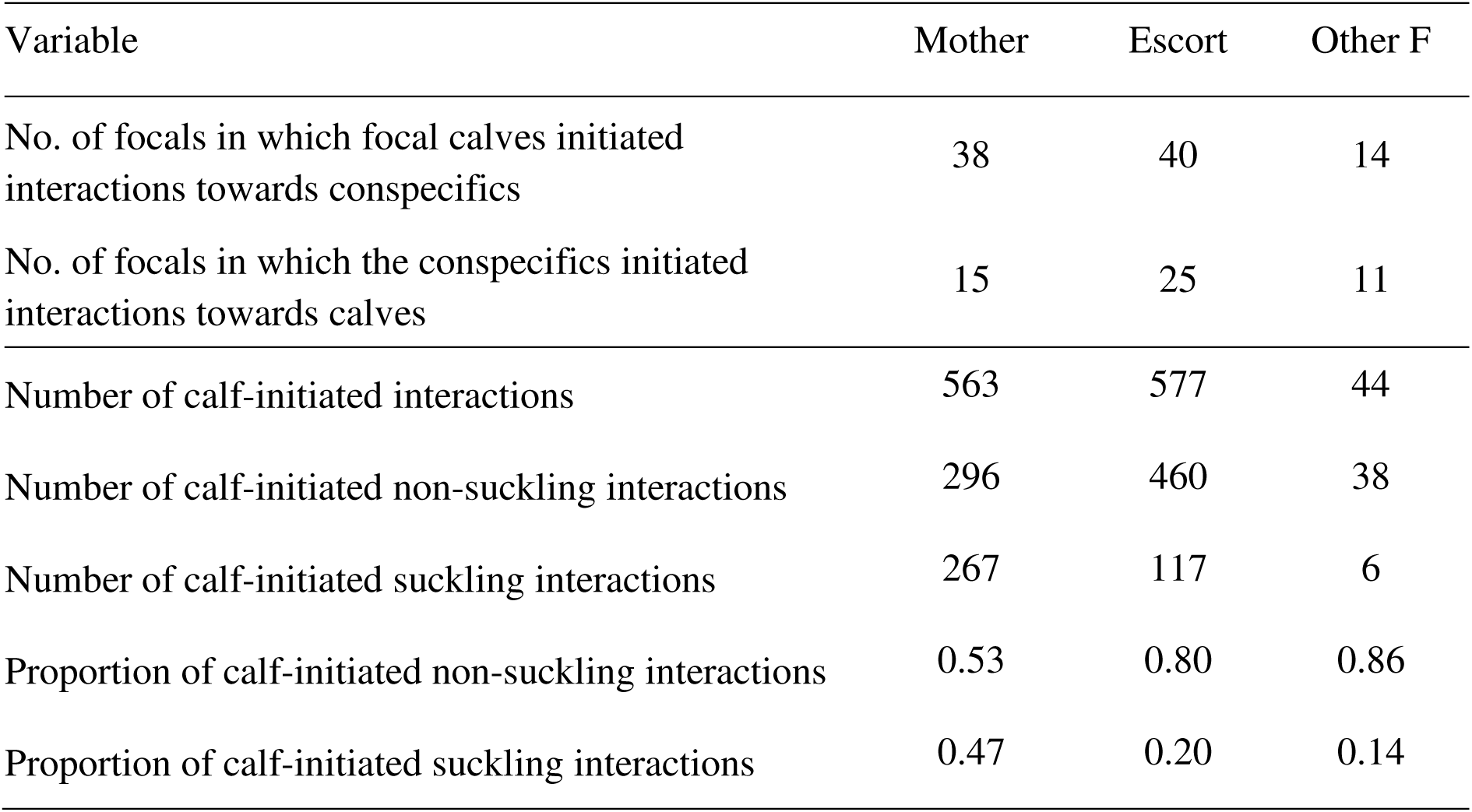
Numbers of focals (out of 40) in which calves and conspecific females initiated interactions towards each other, and the numbers of all, non-suckling, and suckling interactions initiated by calves towards the three conspecific categories of females in these 40 focals. 74.5% of actual suckling was initiated by calves with the mother, 24.5% was initiated with the escorts, which is much higher than that observed in the Amboseli elephants (3.7% of all suckling bouts with females other than the mother; Lee 1987), and only 1% was initiated with other females.

| Variable | Mother | Escort | Other F |
| --- | --- | --- | --- |
| No. of focals in which focal calves initiated interactions towards conspecifics | 38 | 40 | 14 |
| No. of focals in which the conspecifics initiated interactions towards calves | 15 | 25 | 11 |
| Number of calf-initiated interactions | 563 | 577 | 44 |
| Number of calf-initiated non-suckling interactions | 296 | 460 | 38 |
| Number of calf-initiated suckling interactions | 267 | 117 | 6 |
| Proportion of calf-initiated non-suckling interactions | 0.53 | 0.80 | 0.86 |
| Proportion of calf-initiated suckling interactions | 0.47 | 0.20 | 0.14 |

Supplementary Material 10. Effect of calf sex on the numbers of calf-initiated interactions under different behavioural classes.

As mentioned in the main text (see Methods), we ran a nested ANOVA with the log-transformed numbers of interactions under different behavioural classes as the dependent variable to check the effect of calf sex. As before, we could only test the effect of sex on infant calves due to sample size constraints. We found that the effect of calf sex was not significant (Table 1 below).

**Supplementary Material 10, Table 1.** Results of the ANOVA on the log number of different behavioural classes of calf-initiated interactions towards the three conspecific categories of females using 5 males and 5 female infant calves.

| Effect | Effect<br>(F/R) | *SS* | *df* | *MS* | *F* | *P* |
| --- | --- | --- | --- | --- | --- | --- |
| Calf ID(calf sex) | Random | 7.013 | 8 | 0.877 | 2.106 | **0.043** |
| Calf sex | Fixed | 0.458 | 1 | 0.458 | 0.522 | 0.490 |
| Behavioural-class | Fixed | 27.479 | 2 | 13.740 | 17.371 | **<0.001** |
| Conspecific category | Fixed | 39.014 | 2 | 19.507 | 33.393 | **<0.001** |
| Calf ID(calf sex)*Behavioural-class | Random | 12.655 | 16 | 0.791 | 1.900 | **0.030** |
| Calf ID(calf sex)*Conspecific category | Random | 9.347 | 16 | 0.584 | 1.403 | 0.158 |
| Calf sex*Behavioural-class | Fixed | 0.483 | 2 | 0.242 | 0.305 | 0.741 |
| Calf sex*Conspecific category | Fixed | 0.791 | 2 | 0.395 | 0.677 | 0.522 |
| Behavioural-class*Conspecific category | Fixed | 14.876 | 4 | 3.719 | 9.170 | **<0.001** |
| Calf ID(calf sex)*Behavioural class*Conspecific category | Random | 12.979 | 32 | 0.406 | 0.974 | 0.517 |
| Calf sex*Behavioural-class*Conspecific category | Fixed | 1.132 | 4 | 0.283 | 0.698 | 0.599 |
| Error | — | 37.472 | 90 | 0.416 | — | — |

Note: Inferences should not be drawn about Calf ID or interactions with this factor as each calf was sampled in only two focals.

Supplementary Material 11. Behavioural classes of calf-initiated non-suckling interactions.

Of the calf-initiated interactions, there were a total of 539 feeding interactions (Mother: 328, Escort: 198, Other F: 13) for a total of 92.83 minutes (Mother: 63.15 minutes, Escort: 27.88 minutes, Other F: 1.80 minutes), 89 resting interactions (Mother: 36, Escort: 50, Other F: 3) for a total of 198.28 minutes (Mother: 80.48 minutes, Escort: 113.45 minutes, Other F: 4.35 minutes), and 556 social interactions (Mother: 199, Escort: 329, Other F: 28) for a total of 106.88 minutes (Mother: 34.52 minutes, Escort: 70.35 minutes, Other F: 2.02 minutes).

Of the 539 feeding interactions, 390 were suckling interactions, and a majority of them was with mothers (Supplementary Material 7). As mentioned in the main text (see Methods), we performed a nested ANOVA with only calf-initiated non-suckling interactions of the three behavioural classes with conspecific females and found that the pattern of results was the same as that of all calf-initiated interactions (see Table 2 and Figure 6 in the main text and Table 1 and Figure 1 below). Here again, newborn calves initiated similar numbers of interactions as infant calves towards conspecific females under the three behavioural classes (Table 1, Figure 1 below). Tukey’s HSD tests involving feeding interactions showed a different pattern of results from that seen in the analysis using all calf-initiated interactions. Calves initiated similar numbers of feeding and social interactions (95% CI around difference between means for behavioural class: 0.310; Tukey’s HSD: *P*>0.05, Table 2) towards conspecific females when all the calf-initiated interactions were considered. However, this comparison became significant when only calf-initiated non-suckling interactions were considered (95% CI around difference between means for behavioural class: 0.342; Tukey’s HSD: *P*<0.05), as a majority of the calf-initiated feeding interactions were suckling interactions (see Supplementary Material 7). So, calves initiated a greater number of social interactions than non-suckling feeding interactions towards conspecific females (Figure 6b). Similarly, calves initiated a greater number of feeding interactions towards their mothers than towards other females (95% CI around difference between means for behavioural class x conspecific category: 0.502; *P*<0.05, Figure 6a). However, this comparison became non-significant when only calf-initiated non-suckling feeding interactions were considered (95% CI around difference between means for behavioural class: 0.452; *P*>0.05, Figure 6b), as a majority of the suckling interactions of calves were with their mothers (see Supplementary Material 7).

**Supplementary Material 11, Table 1.** Results of the ANOVA on the log number of non-suckling calf-initiated interactions towards the three conspecific categories of females under different behavioural classes. Significant *P* values are marked in bold.

| Effect | Effect<br>(F/R) | *SS* | *df* | *MS* | *F* | *P* |
| --- | --- | --- | --- | --- | --- | --- |
| Age-class | Fixed | 0.283 | 1 | 0.283 | 0.434 | 0.518 |
| Calf ID(Age-class) | Random | 11.717 | 18 | 0.651 | 1.872 | **0.021** |
| Behavioural-class | Fixed | 53.409 | 2 | 26.704 | 22.801 | **<0.001** |
| Conspecific category | Fixed | 43.610 | 2 | 21.805 | 31.139 | **<0.001** |
| Behavioural-class* Conspecific category | Fixed | 11.796 | 4 | 2.949 | 7.385 | **<0.001** |
| Age-class*Behavioural-class | Fixed | 3.639 | 2 | 1.820 | 1.554 | 0.225 |
| Age-class*Conspecific category | Fixed | 0.390 | 2 | 0.195 | 0.278 | 0.759 |
| Age-class*Behavioural-class*Conspecific category | Fixed | 2.857 | 4 | 0.714 | 1.789 | 0.140 |
| Calf ID(Age-class)*Behavioural-class | Random | 42.162 | 36 | 1.171 | 3.368 | **<0.001** |
| Calf ID(Age-class)*Conspecific category | Random | 25.209 | 36 | 0.700 | 2.013 | **0.002** |
| Calf ID(Age-class)*Behavioural-class*Conspecific category | Random | 28.749 | 72 | 0.399 | 1.148 | 0.232 |
| Error | — | 62.601 | 180 | 0.348 | — | — |

Note: Inferences should not be drawn about Calf ID or interactions with this factor as each calf was sampled in only two focals.

**Supplementary Material 11, Figure 1.**
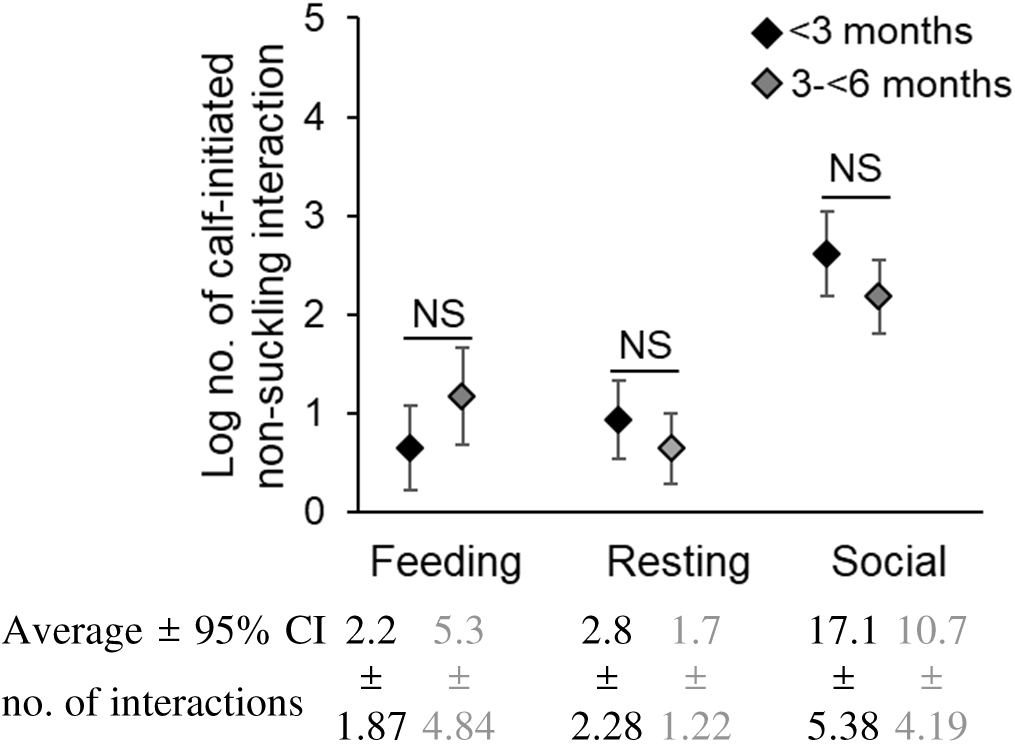
Log numbers of calf-initiated non-suckling interactions of the three behavioural classes by newborn and infant calves towards conspecific females. Error bars are 95% CI. NS indicates lack of statistical significance.

Supplementary Material 12. Termination of calf-initiated interactions.

### All interactions

**Supplementary Material 12, Table 1.** Results of the nested ANOVA on the logit proportion of calf-initiated interactions towards mothers and escorts that were terminated by these females. The asterisks in the *P* values column indicate significance in the ANOVA on the logit proportion of calf-initiated non-suckling interactions (see Supplementary Material 12, Table 2) for comparison.

| Effect | Effect (F/R) | SS | df | MS | F | P |
| --- | --- | --- | --- | --- | --- | --- |
| Calf ID(Age-class) | Random | 1.121 | 14 | 0.080 | 1.387 | 0.216* |
| Age-class | Fixed | 0.236 | 1 | 0.236 | 2.953 | 0.108 |
| Conspecific category | Fixed | 0.036 | 1 | 0.036 | 1.314 | 0.271 |
| Age-class*Conspecific category | Fixed | 0.012 | 1 | 0.012 | 0.440 | 0.518 |
| Calf ID(Age-class)*Conspecific category | Random | 0.388 | 14 | 0.028 | 0.480 | 0.927 |
| Error | — | 1.848 | 32 | 0.058 | — | — |

### Non-suckling interactions

Of the 794 calf-initiated non-suckling interactions with mothers, escorts, and other females, calves terminated 563 interactions, and conspecifics terminated 231. There was no difference between newborn and infant calves in the logit proportions of their non-suckling interactions terminated by their mothers and escorts (Table 1 below). Mothers and escorts terminated similar proportions of calf-initiated non-suckling interactions (Table 1, Figure 1 below). There were insufficient data to examine the effect of calf sex on terminations.

**Supplementary Material 12, Table 2.** Results of the nested ANOVA on the logit proportion of calf-initiated non-suckling interactions towards mothers and escorts that were terminated by these conspecific females. Significant *P* values are marked in bold.

| Effect | Effect (F/R) | SS | df | MS | F | P |
| --- | --- | --- | --- | --- | --- | --- |
| Calf ID(Age-class) | Random | 156.403 | 12 | 13.034 | 2.986 | **0.008** |
| Age-class | Fixed | 50.021 | 1 | 50.021 | 3.838 | 0.074 |
| Conspecific category | Fixed | 7.087 | 1 | 7.087 | 0.831 | 0.380 |
| Age-class*Conspecific category | Fixed | 4.301 | 1 | 4.301 | 0.504 | 0.491 |
| Calf ID(Age-class)*Conspecific category | Random | 102.316 | 12 | 8.526 | 1.954 | 0.071 |
| Error | — | 122.205 | 28 | 4.364 | — | — |

Note: Inferences should not be drawn about Calf ID or interactions with this factor as each calf was sampled in only two focals.

**Supplementary Material 12, Figure 1.**
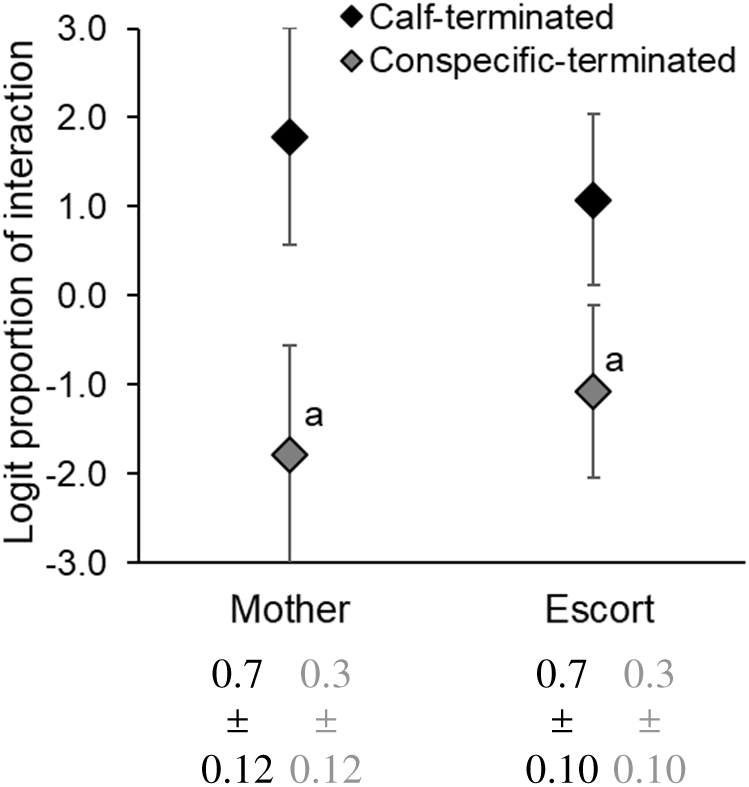
Logit proportions of calf-initiated non-suckling interactions towards their mothers and escorts that were terminated by the calves and by the conspecific females. Error bars are 95% CI. Letters above the data points indicate pattern of statistical significance. Proportions of calf-initiated non-suckling interactions terminated by calves and conspecific females are written below the graph (average ± 95% CI).

Supplementary Material 13. Types of responses from conspecific females for calf-initiated interactions.

### All interactions

**Supplementary Material 13, Table 1.** Results of the ANOVA on the logit proportions of all calf-initiated interactions towards mothers and escorts that elicited a positive or a neutral response. Significant *P* values are marked in bold. The asterisks in the *P* values column indicate significance in the ANOVA on the log numbers of calf-initiated non-suckling interactions (see Supplementary Material 13, Table 2) for comparison.

| Effect | Effect (F/R) | SS | df | MS | F | P |
| --- | --- | --- | --- | --- | --- | --- |
| Calf ID(Age-class) | Random | 17.477 | 14 | 1.248 | 0.241 | 0.997 |
| Age class | Fixed | 1.363 | 1 | 1.363 | 1.092 | 0.314 |
| Response type | Fixed | 766.890 | 1 | 766.890 | 43.888 | **<0.001\*** |
| Conspecific category | Fixed | 0.002 | 1 | 0.002 | 0.002 | 0.966 |
| Calf ID(Age-class)*Response type | Random | 244.635 | 14 | 17.474 | 3.374 | **<0.001\*** |
| Calf ID(Age-class)*Conspecific category | Random | 13.358 | 14 | 0.954 | 0.184 | 0.999 |
| Age class*Response type | Fixed | 76.427 | 1 | 76.427 | 4.374 | 0.055 |
| Age class*Conspecific category | Fixed | 0.273 | 1 | 0.273 | 0.286 | 0.601 |
| Response type*Conspecific category | Fixed | 6.574 | 1 | 6.574 | 1.117 | 0.298 |
| Age class*Response type*Conspecific category | Fixed | 1.882 | 1 | 1.882 | 0.320 | 0.581 |
| Calf ID(Age-class)*Response type*Conspecific category | Random | 82.386 | 14 | 5.885 | 1.136 | 0.345 |
| Error | — | 331.429 | 64 | 5.179 | — | — |

### Non-suckling interactions

After removing the suckling interactions, of the 756 calf-initiated interactions towards mothers and escorts (Supplementary Material 7), there was a positive or a negative response for 174 of the interactions. Similar to the analysis using the full data, we found that there was a significant effect of response type and the interaction between calf ID and response type (Table 2 below, Figure 1 below). Mothers (0.36% of the 563 calf-initiated interactions towards mothers) and escorts (1.39% of the 577 calf-initiated interactions towards escorts) showed a much smaller proportion of aggressive responses than other females (15.91% of the 44 calf-initiated interactions towards other females) towards the calf-initiated interactions.

None of the other effects was significant.

**Supplementary Material 13, Table 2.** Results of the ANOVA on the logit proportions of non-suckling calf-initiated interactions towards mothers and escorts that elicited a positive or a neutral response. Significant *P* values are marked in bold.

| Effect | Effect (F/R) | *SS* | *df* | *MS* | *F* | *P* |
| --- | --- | --- | --- | --- | --- | --- |
| Calf ID(Age-class) | Random | 41.365 | 12 | 3.447 | 0.600 | 0.833 |
| Age class | Fixed | 9.395 | 1 | 9.395 | 2.726 | 0.125 |
| Response type | Fixed | 784.004 | 1 | 784.004 | 40.673 | **<0.001** |
| Conspecific category | Fixed | 1.993 | 1 | 1.993 | 0.888 | 0.365 |
| Calf ID(Age-class)*Response type | Random | 231.309 | 12 | 19.276 | 3.355 | **0.001** |
| Calf ID(Age-class)*Conspecific category | Random | 26.932 | 12 | 2.244 | 0.391 | 0.961 |
| Age class*Response type | Fixed | 74.367 | 1 | 74.367 | 3.858 | 0.073 |
| Age class*Conspecific category | Fixed | 3.073 | 1 | 3.073 | 1.369 | 0.265 |
| Response type*Conspecific category | Fixed | 10.609 | 1 | 10.609 | 1.136 | 0.294 |
| Age class*Response type*Conspecific category | Fixed | 1.880 | 1 | 1.880 | 0.201 | 0.662 |
| Calf ID(Age-class)*Response type*Conspecific category | Random | 112.050 | 12 | 9.338 | 1.625 | 0.111 |
| Error | — | 321.774 | 56 | 5.746 | — | — |

Note: Inferences should not be drawn about Calf ID or interactions with this factor as each calf was sampled in only two focals.

**Supplementary Material 13, Figure 1.**
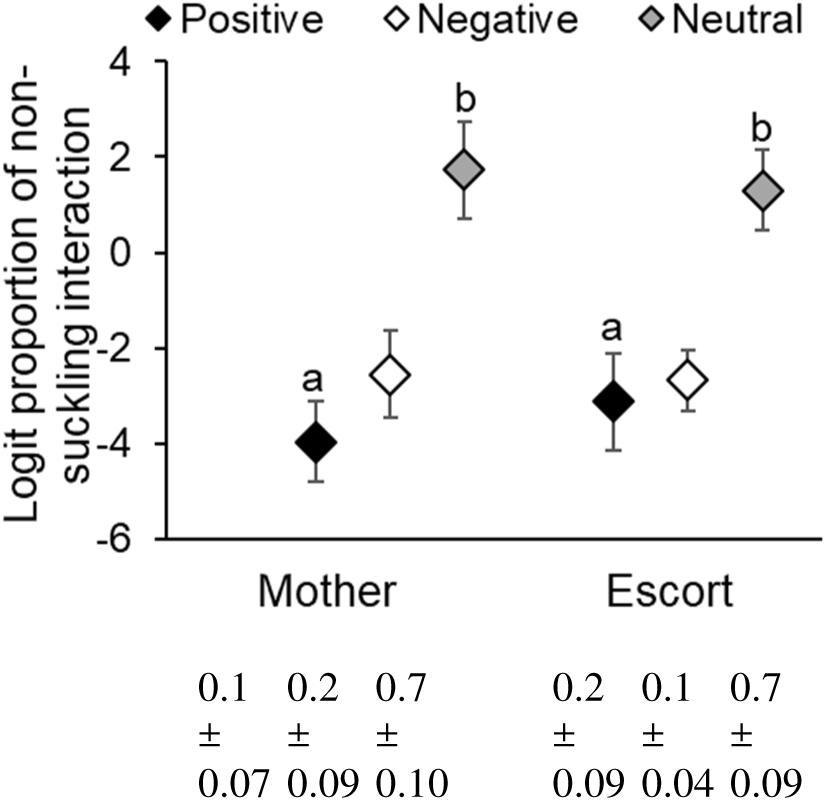
Logit proportions of calf-initiated non-suckling interactions towards mothers and escorts that elicited a positive, neutral, and negative responses from the conspecific females. Error bars are 95% CI. Letters above the data points indicate patterns of statistical significance (a<b). Proportions of calf-initiated non-suckling interactions that elicited a positive, neutral, and negative responses from the conspecific females are written below the graph (average ± 95% CI).

